# Decoding Gene Responsiveness to Synthetic Chromatin Reader-Actuators with Multi-Modal Epigenomic Profiling

**DOI:** 10.1101/2025.11.26.690833

**Authors:** Seong Hu Kim, Isioma Enwerem-Lackland, Natecia L. Williams, Rachel Fisher, Christopher Plaisier, Karmella A. Haynes

**Affiliations:** Wallace H. Coulter Department of Biomedical Engineering, Emory University, Atlanta, GA; School of Biological and Health Systems Engineering, Ira A. Fulton School of Engineering, Arizona State University, Tempe, AZ; John Schufeldt School of Medicine and Medical Engineering, Arizona State University, Tempe, AZ

**Keywords:** epigenome engineering, polycomb, chromatin, gene regulation, topologically associating domains, machine learning, breast cancer

## Abstract

Cell identity is regulated by chromatin states that encode gene regulatory memory and shape responses to new inputs. To investigate how chromatin context influences inducibility in differentiated cells, we employed engineered synthetic reader-actuators (SRAs), fusion proteins containing the polycomb chromodomain (PCD) that binds H3K27me3. In MCF7 breast cancer cells, we mapped PCD-fusion occupancy by ChIP-seq and used RNA-seq to identify temporally resolved gene activation patterns. ChIP-seq profiling and machine learning models (MLM) demonstrated that PCD-fusion binding was predicted by the presence of H3K27me3, H4K20me1, or H3K36me3 enrichment, suggesting selective accessibility at enhancers and chromatin transition zones. Among genes with PCD-fusion enriched enhancers, the SRA-induced subset was distinguished by promoter features including bivalent histone modifications and enrichment of transcriptional repressors REST and MTA1. Collectively, our results demonstrate that SRA responsiveness depends on a specific chromatin signature beyond H3K27me3 alone. This study demonstrates the power of SRAs to dissect inducible chromatin features in their native genomic context, and suggests that epigenetically repressed regions in differentiated cells can retain regulatory plasticity.

## INTRODUCTION

A promising approach for cell engineering is to leverage the structural plasticity of chromatin to alter gene transcription profiles and change key phenotypic features that determine cell function to ultimately reshape cell states.^1,2^ Histone protein octamers form nucleosome complexes with genomic DNA, where each nucleosome is spaced by nearly zero base pairs in compact chromatin, or several hundred base pairs in less condensed chromatin.^3^ The histones are subjected to an array of post-translational modifications, resulting in an extensive layer of biochemical information that impacts transcriptional activity for nearly all genes. One major challenge for controlling chromatin is to determine the minimum set of genomic or epigenomic features needed to predict which genes will respond to specific chromatin perturbations. Achieving this goal would enable rational control over multi-gene networks relevant to development, regeneration, and disease.

Studies of dynamic chromatin states at regulatory elements in differentiating stem cells provide insights that inform strategies to engineer cell identity by modulating chromatin architecture. Non-coding sequences called enhancers maintain a chromatin-encoded poised “bivalent” state marked by H3K4me1/2 and H3K27me3, which allows rapid and lineage-specific induction of developmental genes upon differentiation cues.^4–9^ Before activation, enhancers enter a primed state marked by H3K4me1, with or without H3K27me3.^7,10^ Upon activation, enhancers retain H3K4me1, lose H3K27me3, gain H3K27ac, and often form physical contacts with the promoter near a gene’s transcriptional start site (TSS).^11–14^ Enhancers can also become inactivated or “decommissioned” by losing H3K4me1, gaining DNA methylation, or by reinforcing polycomb protein complexes.^15,16^ Even after differentiation, many enhancers maintain a poised or primed state, allowing gene expression reprogramming.^17^ A second class of regulatory elements, polycomb response elements (PREs), can also switch states, but unlike enhancers, PREs are thought to support “memory” through a stable chromatin configuration. Initially characterized in *Drosophila*, mammalian PREs lack conserved DNA sequences.^18,19^ Their identification required functional mapping approaches, which ultimately identified PREs at the murine *MafB* gene,^20^ and in the human *HOX* clusters and other loci.^21–24^

A central feature of regulator states is H3K27me3, which is generated by the EZH2 subunit in polycomb repressive complex 2 (PRC2). Specifically targeting H3K27me3 could offer an approach to switch on silent genes in a chromatin context-dependent manner. State-of-the-art methods for altering chromatin states rely on loss-of-function perturbations, such as genetic knock-down, small molecule inhibitors, or CRISPR-based knockouts of chromatin proteins.^25,26^ While these tools are powerful, they are limited in several respects. First, they can fail to activate genes in cells lacking functional transcriptional cofactors. Furthermore, they affect both nuclear and cytoplasmic roles of chromatin-associated enzymes. Finally, they do not easily resolve temporal dynamics of gene activation. These limitations motivate the need for gain-of-function tools that directly test the functional role of chromatin features across the genome.

Fusion protein technologies can address the limitations of traditional loss-of-function approaches by enabling control over chromatin activity. For example, epigenome editing using dCas9-based chromatin modifiers^27^ and endonuclease-mediated excision of regulatory regions^28^ can target specific loci while minimizing off-target effects through nuclear localization signals and sequence-specific recruitment. However, these methods are typically limited to one or a few genomic sites at a time. In contrast, synthetic reader-actuators (SRAs) offer a scalable, genome-wide approach. SRAs are engineered from natural regulators that bind histone post-translational modifications (PTMs) or DNA methylation, and consist of a chromatin-binding domain fused to a transcriptional effector domain, retaining core functions while minimizing size and complexity.^29,30^ In our previous work, we fused the H3K27me3-binding domain of CBX8 (PCD) to the transcriptional activator VP64 and expressed this fusion in various cancer cell lines. Transcriptomic analysis showed that the SRA induced widespread activation of silent or lowly expressed genes, but many H3K27me3-enriched sites remained unperturbed by SRA activity. Therefore, a key open question remains: what additional chromatin features determine which H3K27me3-enriched loci are switchable in response to targeted activators?

Here, we address this question by combining ChIP-seq of a fluorescent protein-tagged PCD with time-resolved RNA-seq after direct perturbation of chromatin via SRAs in transgenic MCF7 breast cancer cells. The genomic distribution of both PCD-fusion proteins showed enrichment in intronic and distal intergenic regions, sites that are generally associated with enhancer elements that contain H3K4me1, and H3K27me3 or H3K27ac in poised or active states respectively. Histone PTM overlap analyses at PCD-fusion binding sites identified H3K27me3 and H3K4me1 as significant marks, as well as H3K36me3 and H4K20me1, features associated with transcriptional transition zones. Perturbation of these sites with SRA revealed a transcriptionally responsive set of 397 genes, including many known polycomb protein targets such as *HOXB* and *HOXC* loci, and *MAFA*. Most showed rapid and transient activation over 10 - 12 hours. Machine learning modeling (MLM) revealed a combinatorial histone profile for PCD-fusion binding sites, and that single marks are not sufficient to predict binding. MLM also indicated that SRA-mediated activation is associated with a low baseline gene expression level, a bivalent histone PTM signature and repressors REST and MTA1 at promoters, and the H3K36me3 reader protein HDGF at enhancers. This work provides a framework for predicting the outcomes of targeting H3K27me3 based on a minimal set of feature annotations.

## RESULTS

### PCD-fusion proteins map to intronic and intergenic loci enriched for poised and transitional regulatory marks

First, we tested the hypothesis that fusion proteins carrying the polycomb chromodomain (PCD), derived from the CBX8 protein, would be observed at sites in the genome that are enriched for H3K27me3. We carried out ChIP-seq using crosslinked chromatin from MCF7 cells that expressed PCD tagged with an mCherry red fluorescent protein (RFP), from a stably-inserted sleeping beauty transgene (pSBtetTA-GP) with a doxycycline-inducible promoter (**Figure 1A**). Another cell line was engineered to express the same fusion, with the addition of a transcriptional activation domain, PCD-RFP-VP64 (synthetic reader-actuator, SRA), as we have previously reported.^31^ We verified intact transgene inserts via PCR of genomic DNA with transgene-specific primers, followed by Sanger sequencing (**Figure 1B**). Sub-cloning of a DpnII-digested fragment library and transgene specific PCR (as previously described^32^) was used to map the genomic location of each insert at chromosome 6. The SRA transgene is located in the intron of *RSPH9*, while PCD-RFP is located in non-coding intergenic DNA (**Figure 1C**). Fluorescence microscopy of cells treated with 1.0 µg/mL doxycycline confirmed expression and nuclear localization of the RFP-tagged chromatin proteins (**Figure 1D**). RT-qPCR confirmed that expression of both the SRA and PCD-RFP transgenes was induced by doxycycline in a dose-dependent manner (**Figure 1E**).

**Figure 1.**
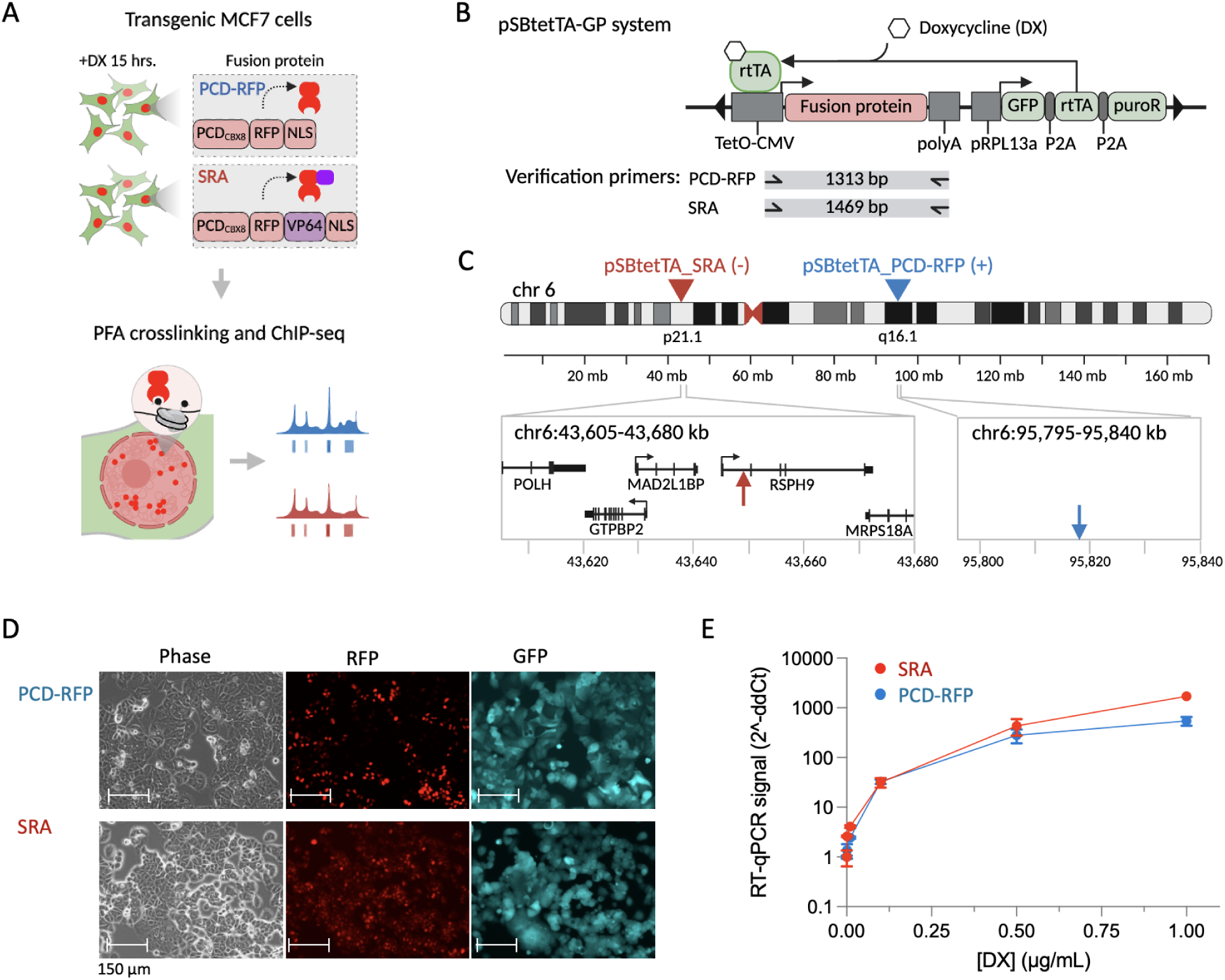
Experimental design and validation of transgenes in MCF7 cells. (A) Experimental strategy for the ChIP-seq experiment. (B) Map of the pSBtetTA-GP transgene cassette and PCR primers used to amplify the transgenes from genomic DNA. (C) The pSBtet transgene inserts were mapped to chromosome 6. (D) Representative images of 1.0 µg/mL doxycycline-treated transgenic MCF7 cells. (E) Measurement of transgene expression levels via RT-qPCR with universal primers for mCherry. Values were normalized by RT-qPCR of GAPDH.

PCD-RFP-expressing cells were treated with 1.0 µg/mL doxycycline and crosslinked with 1% formaldehyde after 15 hours to capture early fusion protein binding events. PCD-RFP ChIP-seq signals showed broad distribution patterns (**Figure 2A**) and peaks appeared over large H3K27me3-enriched domains (**Figure 2B**), as expected for PCD binding activity.^33^ We carried out the same experiments for SRA-expressing cells treated with titrating concentrations of doxycycline (0.0, 0.1, 0.5, and 1.0 µg/mL) since SRA over-expression significantly alters the transcription profile and potentially the phenotype of the cells.^31^ SRA ChIP-seq signals increased with higher doxycycline dosage and peaks generally tracked with PCD-RFP binding (**Figure** 2A, B). The higher abundance of PCD-RFP peaks compared to SRA might be due to a greater read-depth (>6x) in this smaller sample set.

**Figure 2.**
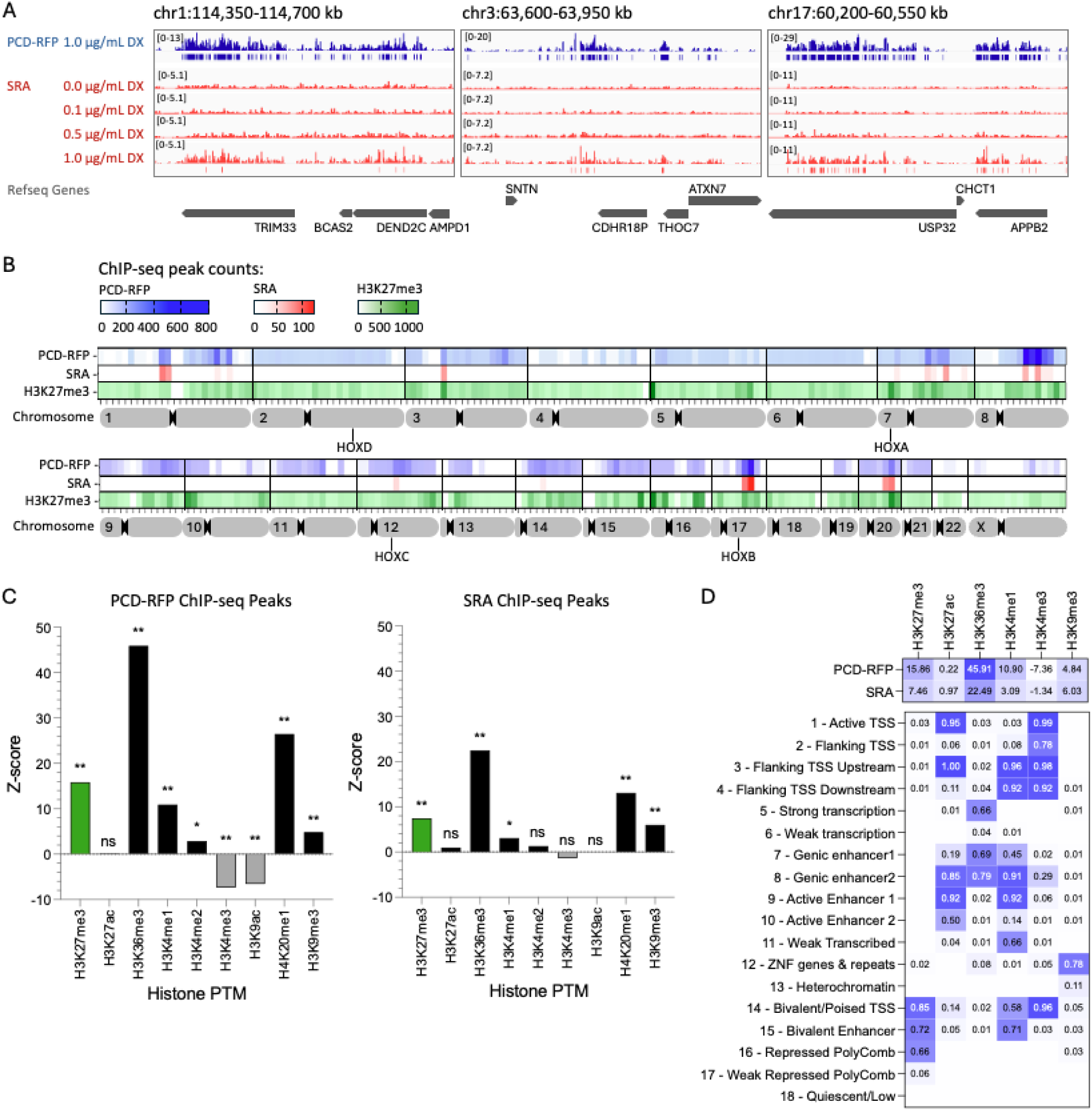
Genomic distributions and chromatin feature overlaps for SRA and PCD-RFP ChIP-seq peaks. (A) Representative regions of SRA and PCD-RFP enrichment. Bed and bigWig file tracks were visualized using Integrative Genomics Viewer (IGV).^34^ (B) Heat maps show ChIP-seq peak counts in 10 Mb intervals across the human genome (GRCh38/hg38). (C) Z-scores for computed overlaps of chromatin features with ChIP-seq peak regions for PCD-RFP and SRA. *p*-value = **0.001, *0.004, ns = not significant. (D) Top table: Z-score values of histone PTM ChIP-seq peaks and PCD-RFP or SRA ChIP-seq peak overlaps. Bottom table: histone PTM enrichment values for the MCF7 18-state Chrom HMM model from the Epigenome Roadmap.^35^ Values for the general 15-state model^36^ are presented in Supplemental Figure S1.

Polycomb response elements (PRE) represent evolutionarily conserved hubs of PRC-mediated gene regulation, and are often enriched for H3K27me3. The region with the highest frequency of PCD-fusion ChIP-seq peaks near a known PRE is at chromosome 17, where the HOXB cluster is located at ∼48.5 - 48.6 Mb. We observed a few PCD-fusion peaks near HOX loci on chromosome 12 roughly 1 Mb from the HOXC cluster (∼53.9 - 54.1 Mb), and on chromosome 7 about 1.4 Mb from HOXA (chr7, ∼27.0 - 27.2 Mb). We did not observe high frequencies of PCD-fusion peaks near HOXD (chr2, ∼176.0 - 176.2 Mb) regions.

To determine chromatin context at all PCD-RFP and SRA binding sites, we computed Z-scores for the overlap between ChIP-seq peaks and histone PTM datasets for MCF7 cells (**Figure 2C**). PCD-RFP peak regions showed the strongest enrichment for H3K36me3 (Z = 45.9), H4K20me1 (Z = 26.5), and H3K27me3 (Z = 15.9), with additional enrichment for H3K4me1/2 and H3K9me3. Conversely, PCD-RFP regions were significantly depleted for promoter-associated marks H3K4me3 (Z = -7.4) and H3K9ac (Z = -6.6), suggesting exclusion from active promoters. SRA binding regions showed a similar but more restricted pattern, with enrichment for H3K36me3 (Z = 22.5), H4K20me1 (Z = 13.1), H3K27me3 (Z = 7.5), H3K9me3 (Z = 6.0), and H3K4me1 (Z = 3.1). An alternative analysis using fold enrichment of multiple histone ChIP-seq datasets (ChIP-Atlas fold enrichment tool) was generally consistent with these results (**Supplemental Figure S1**).

To better understand the gene regulatory states associated with PCD-fusion binding sites, we compared the overlap profiles to regulatory states defined by histone PTM combinations (ChromHMM and Roadmap Epigenomics Consortium).^36,37^ Comparisons with a general 15-state model, and an MCF7 18-state model revealed that PCD-RFP and SRA binding sites are enriched at regions associated with transcriptionally active or poised chromatin (**Figure 2D, Supplemental Figure S1**). Higher values for H3K36me3, H3K4me1, H4K20me1, and H3K27me3, correspond to genic enhancers, bivalent enhancers, and actively transcribed gene bodies. The significant depletion of promoter-associated marks such as H3K4me3 and H3K9ac at both PCD-RFP and SRA sites supports their exclusion from canonical promoter regions and highlights binding to regulatory elements embedded within repressive or transitional chromatin. These results are consistent with previously reported mapping of endogenous CBX8 in mouse embryonic stem cells.^38^ Therefore, accessibility of H3K27me3 to the PCD domain may be determined by the physical chromatin structure collectively formed by other histone PTMs.

Annotation of ChIP-seq peaks using ChIPseeker revealed that PCD-RFP and SRA predominantly localize to intronic and distal intergenic regions (Figure 3A, B). The majority of PCD-RFP-bound regions overlapped annotated introns in genic regions, followed by distal intergenic regions. A similar distribution with smaller intersection sizes was observed for SRA, reflecting a narrower but consistent binding pattern. These findings align with the chromatin feature enrichment analyses, where both fusions were associated with histone modifications indicative of transcriptionally poised or structurally dynamic chromatin, such as H3K36me3, H3K4me1, H4K20me1, and H3K27me3. The enrichment in introns and distal intergenic regions suggests engagement with non-promoter regulatory elements, including poised enhancers, gene body-associated regulatory elements, or transcriptional transition zones.

**Figure 3.**
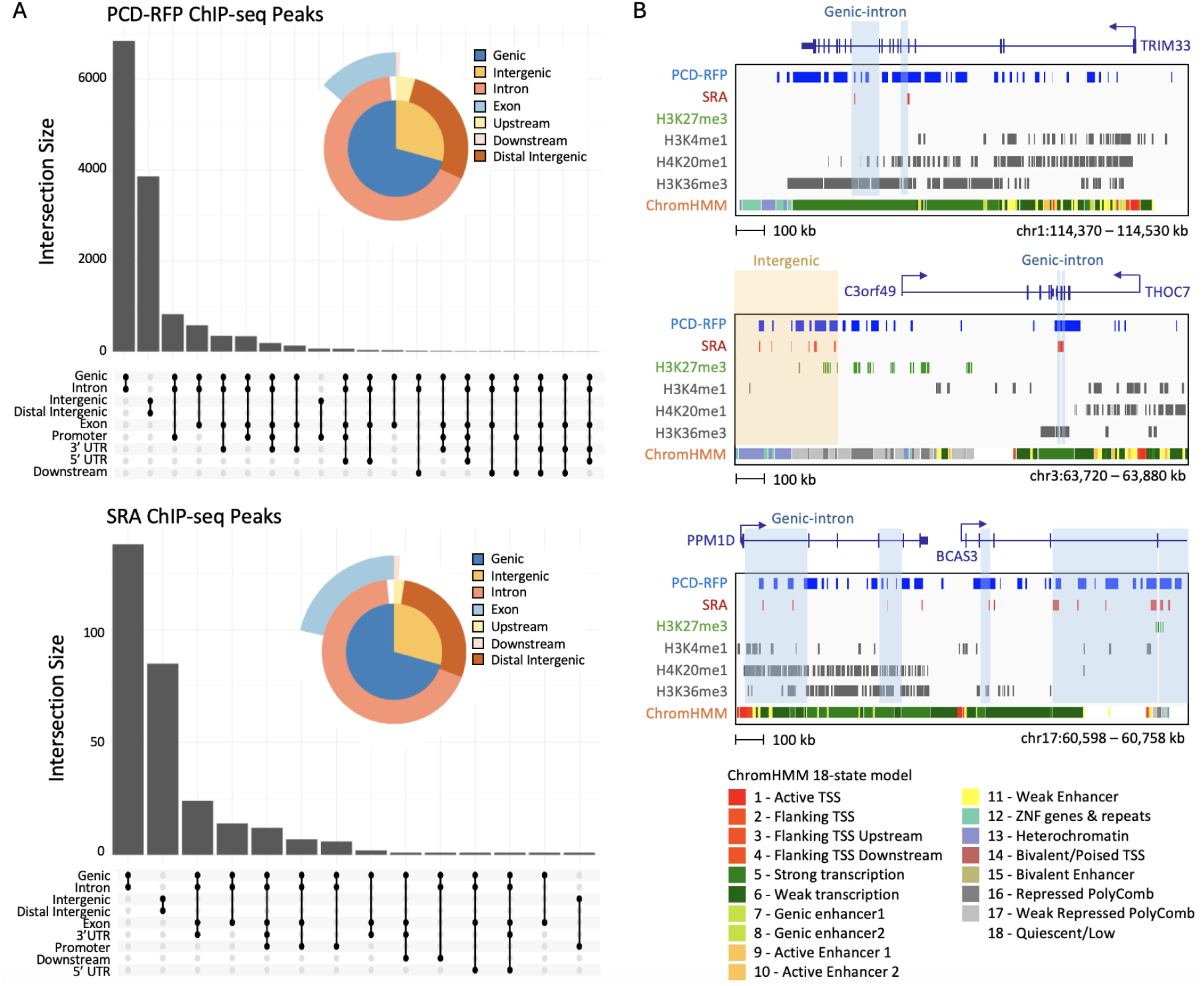
Annotations of regions with PCD-fusion ChIP-seq peaks. (A) ChIPseeker (Bioconductor) was used to compute peak annotations. (B) Representative genic-intron and distal intergenic regions containing ChIP-seq peaks for SRA and PCD-RFP. Genic-intron and distal intergenic regions containing both PCD-RFP and SRA ChIP-seq peaks are shaded.

### Machine learning reveals that combined histone PTM features are required to predict PCD-RFP binding sites in native chromatin

To quantitatively determine the set of chromatin features that predict PCD-RFP binding, an XGBoost machine learning model was trained using 9 histone PTMs (H3K4me1, H3K4me2, H3K4me3, H3K9ac, H3K9me3, H3K27ac, H3K27me3, H3K36me3, H4K20me1) as input features. The model using all 9 features achieved an area under the receiver operating characteristic (AUROC) of 0.96 (**Figure 4A**). The model reached an area under the precision-recall curve (AUCPR) of 0.68, with the random baseline of 0.041 (**Figure 4B**). We used SHAP (SHapley Additive exPlanations) to interpret the model and rank the importance of each histone mark. The feature ranking bar chart (**Figure 4C**) identified H3K4me3, H3K9ac, and H4K20me1 as the three most impactful features for the model’s predictions. The SHAP bee swarm plot (**Figure 4D**) revealed that high values of H3K4me3, H3K9ac, and H3K27ac were clustered at negative SHAP values, which corresponds to the absence of PCD-RFP binding. Conversely, high values of H4K20me1, H3K36me3, H3K9me3, and H3K4me1 were clustered at positive SHAP values, indicating the presence of PCD-RFP binding. H3K27me3 showed a bipolar predictive power with high values clustered at both negative and positive SHAP values, consistent with the idea that PCD-fusions bind a select sub-set of H3K27me3 sites. Finally, we compared the full 9-histone PTM model to simpler models trained on only a single histone PTM (**Figure 4E**). While the H3K4me3-only model achieved the highest single-feature AUROC (0.88) and AUCPR (0.48), the values were lower than those of the full 9-histone PTM model.

**Figure 4.**
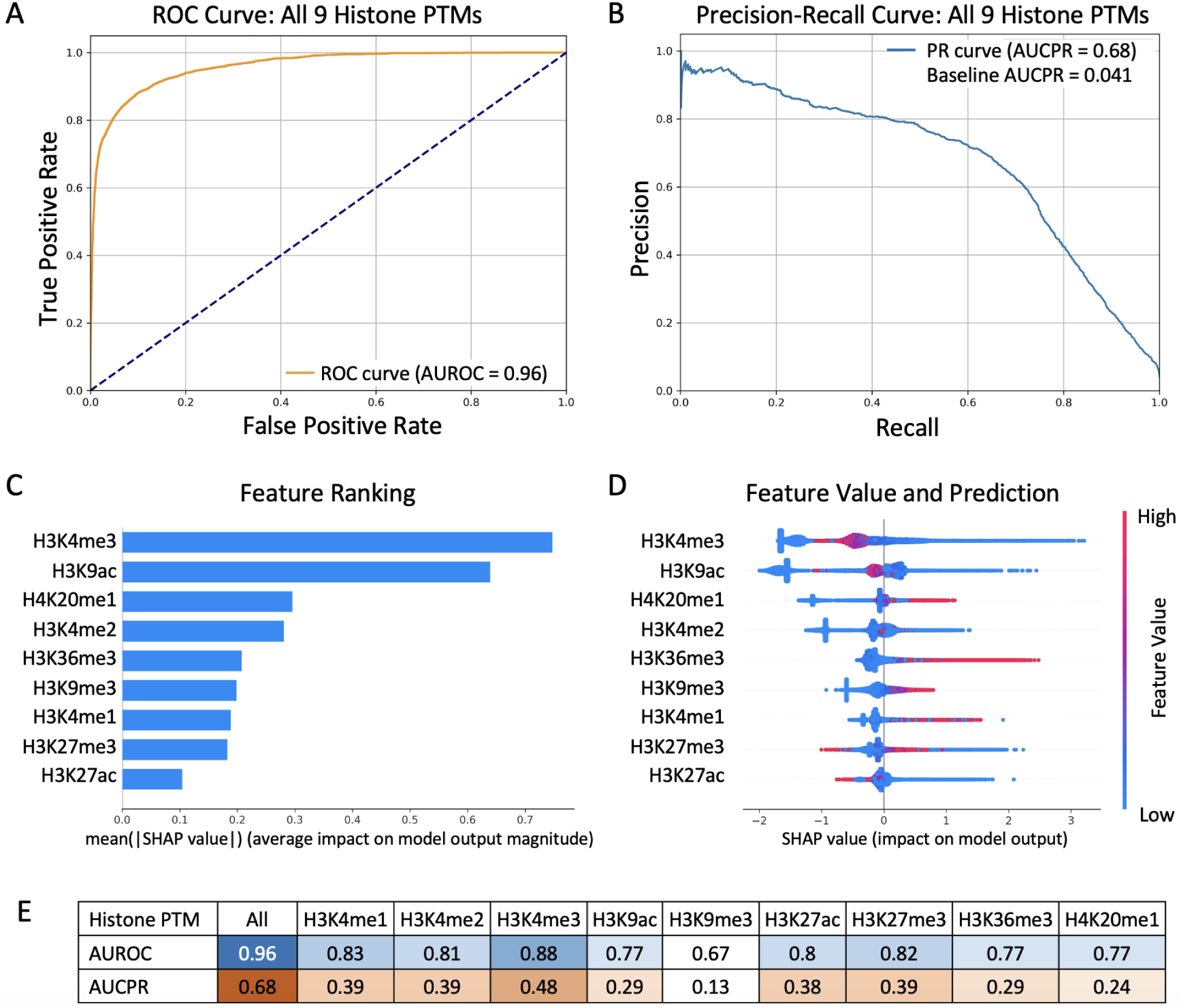
PCD-RFP genome-wide binding prediction using XGBoost. (A) ROC curve and (B) PR curve for predicting PCD-RFP binding using all 9 histone PTMs. (C) SHAP bar chart ranking feature importance for PCD-RFP binding prediction. (D) SHAP bee swarm chart indicating distribution of each feature values respective to SHAP values. (E) AUROC and AUCPR values when predicting PCD-RFP binding using all 9 histone PTMs or only using a single indicated histone PTM.

### Epigenetic perturbation with SRA reveals transcriptionally-responsive topologically associating domains (TADs)

Given the possibility that the PCD-fusion proteins target regulatory elements, we asked whether potential target genes could be upregulated by SRA, which carries the VP64 transcriptional activation domain. We performed time-course RNA-seq by collecting samples at 10, 24, and 48 hours after induction with 1.0 μg/mL doxycycline, compared to non-expressing control cells (**Figure 5A**). Cells expressing a RFP-VP64 fusion that lacks the PCD and cells expressing PCD-RFP were included as negative controls. Time-course profiles of genes upregulated in SRA-expressing cells stratified into 9 classes, suggesting distinct dynamic transcriptional responses. We then categorized 397 SRA-specific differentially upregulated genes (UpDEGs) into six major dynamic response classes based on time points at which they were activated at least 1.5-fold (log₂FC ≥ 0.58, FDR < 0.05): early transient (158 genes), early sustained (68), late transient (51), late sustained (44), delayed (63), and stochastic (13) (**Figure 5B**).

**Figure 5.**
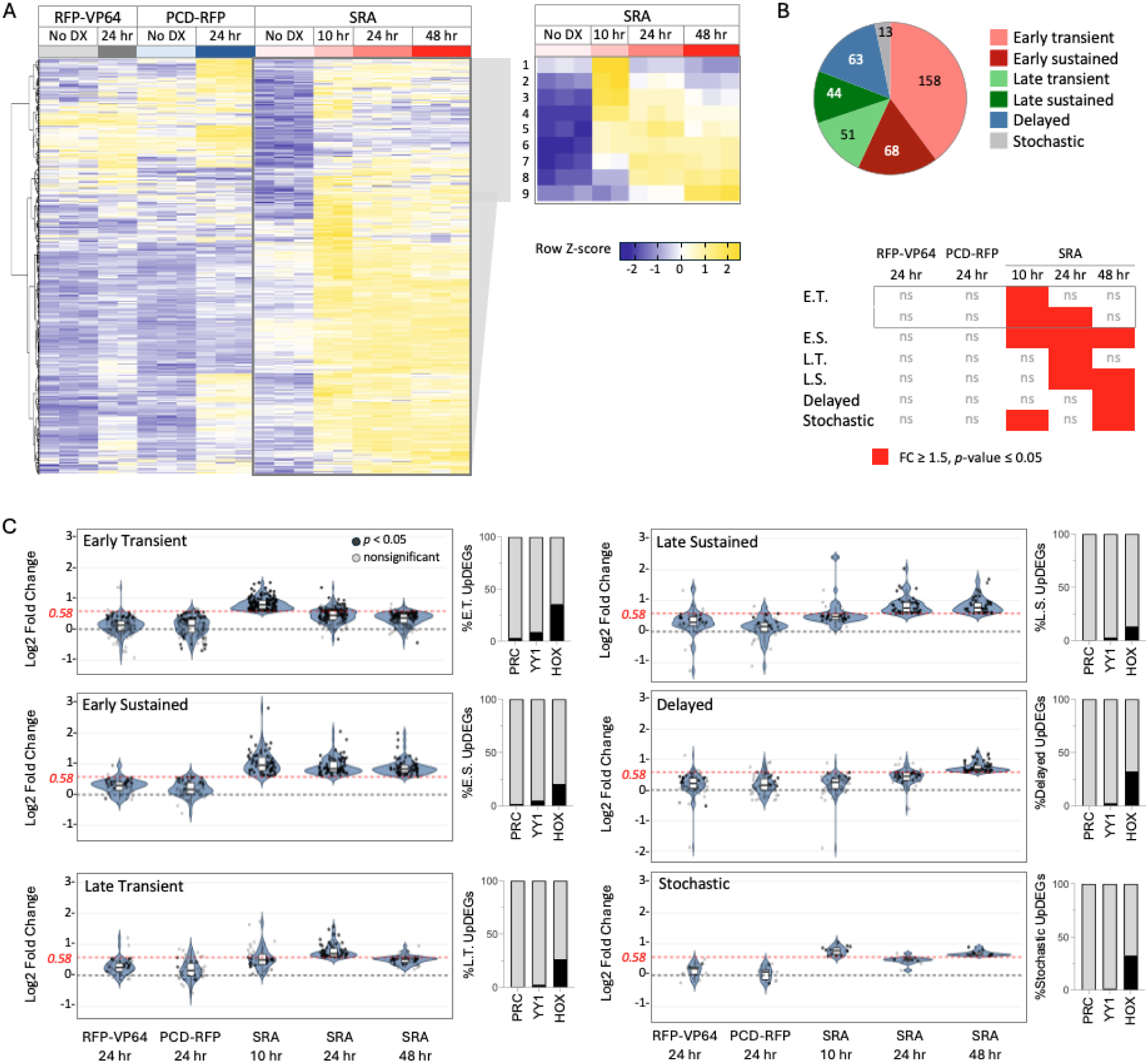
Time-course transcriptome profiling of MCF7 cells with PCD-fusion proteins. (A) The heat map shows normalized RNA-seq values (Z-scores) for triplicate samples for all conditions. Samples identified as outliers were omitted from further analyses. The call-out heatmap shows stratification of genes by Z-scores from SRA-expressing cells. (B) Numbers of SRA-activated UpDEGs in each class are shown in the pie chart. In the table, shaded areas indicate conditions where all genes in each class are significantly upregulated (FC ≥ 1.5, FDR-adjusted *p*-value ≤ 0.05). ns = no significant change observed. (C) Violin plots show log2 fold change RNA-seq values for six dynamic response classes of SRA UpDEGs. Opaque points: *padj* ≤ 0.05; transparent points: *padj* > 0.05 or N/A). Bar charts show the percentage of UpDEGs that were activated after PRC2 knockdown (PRC) in other studies,^39,40^ and have promoters with binding sites for YY1, or at least one upregulated HOX transcription factor. Bar chart gene symbols are listed in **Supplemental Table S1**.

The early sustained genes, representing 17.1% of all UpDEGs, suggest a rapid switch to a stable activated state. This state might be reinforced by transcription factors that are activated by SRA, including HOXB6/9/13 and HOXC8/9/10 (**Supplemental Table S1**). Roughly half of all UpDEGs were early or late transient responders (52.64%) where expression decreased by 48 hours, suggesting that VP64 does not stably overcome the repressed epigenetic state at most activated loci. Late sustained and delayed genes were activated at 24 and 48 hours, or at 48 hours only may represent loci that are less accessible to the SRA, or indirectly regulated by a transcription factor that had been expressed from a different locus at an earlier time point.

Our kinetic classification highlights distinct modes of gene activation over time, suggesting that the SRA unmasks layers of latent transcriptional potential within the epigenome. To determine if different transcriptional regulators underlie these differences, we investigated the frequencies of genes regulated by polycomb proteins, and genes that may be secondary targets of transcription factors encoded by UpDEGs (**Figure 5C**). We identified a few genes within each UpDEG class that corresponded with genes reported to become upregulated by PRC2 knockdown in MCF7 cells in other studies.^39,40^ The regulator YY1, which is associated with polycomb-mediated regulation,^41,42^ has binding sites in 1.3 - 8.9% of UpDEGs from each class. Several master regulators from the HOX family of transcription factors appear in the early transient class of UpDEGs: HOXB6/9/13 and HOXC8/9/10. 13.2 - 35.4% of UpDEGs in every class have binding sites for these transcription factors, suggesting that SRAs may activate networks of genes through HOX transcription factor activity.

In interphase chromatin, polycomb proteins at H3K27me3-marked regions can drive the formation of 3-D loops between enhancers and promoters,^43^ and enhancers are known to regulate genes within physically insulated regions called topologically associating domains (TADs).^44–47^ This information prompted us to analyze the frequency of PCD-fusion binding sites within TADs to gain insight into groups of genes that might be co-regulated through enhancer targeting. Boundaries of TADs were mapped by using insulation scores from MCF7 Hi-C data (Barutcu et al.^48^). The authors excluded TAD boundaries with insulation scores less than 0.15 due to non-reproducibility. We observed maximum co-occupancy between PCD-RFP sites and UpDEGs when we excluded insulation scores less than 0.4. This threshold resulted in 1581 total topologically insulated regions with an average size of ∼2 Mb, about twice the size of a typical 1 Mb TAD (**Figure 6A**). Other work has shown enhancer activity across adjacent TADs in mammalian cells,^49^ therefore it is possible that SRA-mediated activation might not be constrained within single TADs. Within all insulated regions containing UpDEGs, we observed no strong biases in dynamic response classes (**Figure 6A**). Similarly, PCD-RFP binding site-containing regions did not exhibit a clear UpDEG sub-class bias. In several cases, very large PCD-fusion binding-enriched domains (∼20 Mb) overlapped with or were adjacent to regions containing UpDEGs. Other UpDEG regions were PCD-fusion binding site poor (e.g. chr17 ∼30 - 45 Mb). The lack of a very clear intra-TAD relationship between PCD binding sites and UpDEGs raised a critical question: what chromatin or genomic features determine whether a gene is responsive to SRA-mediated activation?

**Figure 6.**
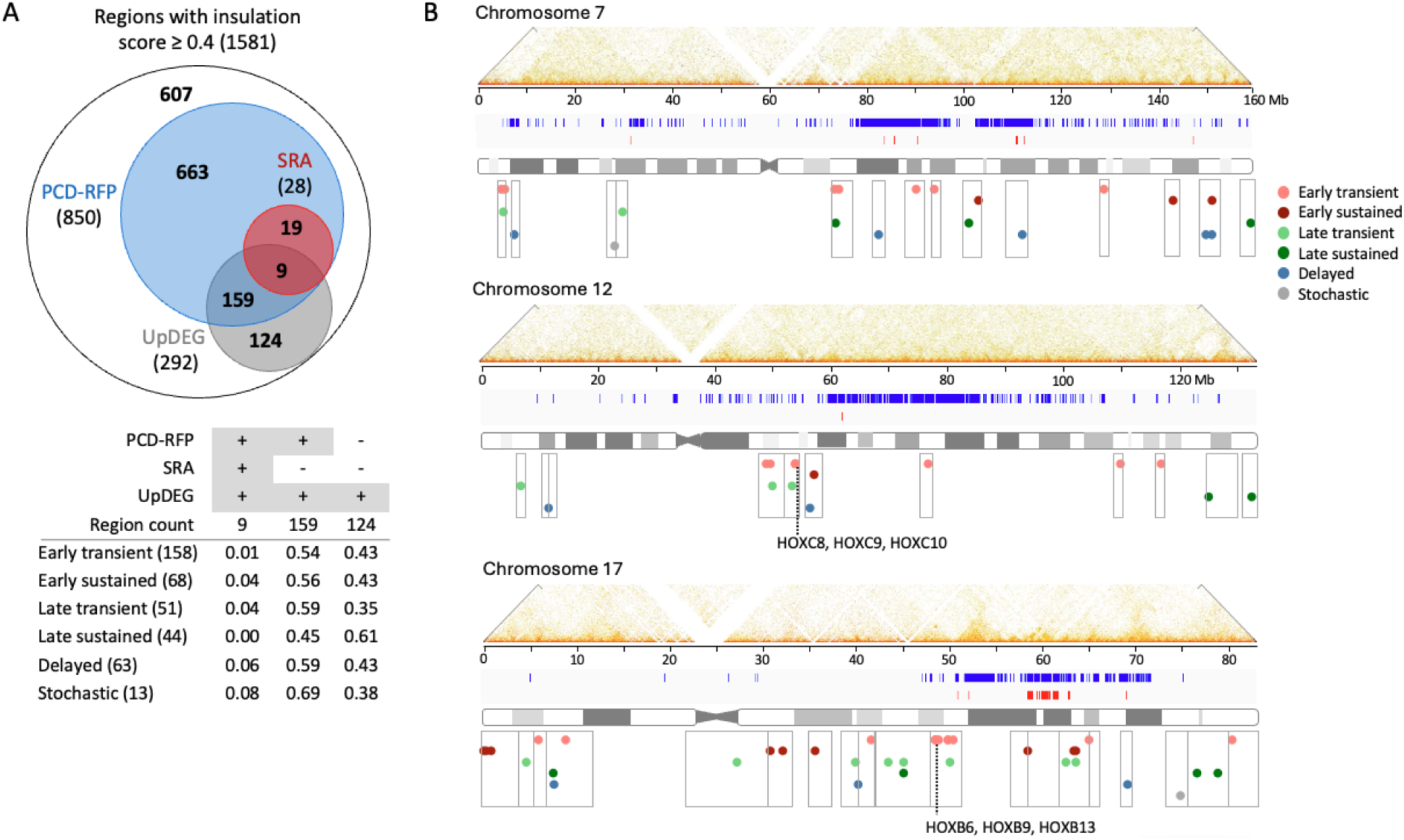
UpDEG and ChIP-seq peak distribution within topologically insulated regions. (A) Venn diagram showing overlap of insulated regions (score ≥ 0.4) categorized by the presence of at least one PCD-RFP or SRA ChIP-seq peak, or SRA UpDEG. The table shows UpDEG totals and frequencies by dynamic class within insulated regions that contain SRA and/or PCD-RFP, and those that do not. (B) Maps for chromosomes 7, 12, and 17 show Hi-C contact heatmaps, tracks for PCD-RFP and SRA ChIP-seq peaks, and transcription start site positions of UpDEGs. Grey boxes mark insulated regions that contain UpDEGs.

### Promoter PCD-RFP enrichment and chromatin state are key predictors of SRA-specific UpDEGs

To identify the regulatory features that distinguish SRA-upregulated genes (UpDEGs) from non-responsive genes (non-DEGs), we developed a multi-omics machine learning pipeline (**Figure 7A**). This model was trained on gene-enhancer pairs (GeneCards Genehancer^50^) where each gene was represented by a single enhancer that showed the maximum PCD-RFP binding signal (**Figure 7B)**. A multi-omic feature matrix was constructed from over 20,000 potential predictors, including 9 histone PTMs, 116 transcription factors, DNase/ATAC-seq signals, and Gene Ontology (GO) terms. These features were mapped to specific genomic regions: the enhancer, core promoter (CP), upstream proximal promoter (USPP), and downstream proximal promoter (DSPP). A multi-stage Recursive Feature Elimination (RFE) process, guided by AUCPR, reduced this to an optimized set of 150 features. The final ensemble model, which averaged the output of XGBoost, LightGBM, CatBoost, and Random Forest, was evaluated on a held-out test set of chromosomes. The model achieved an AUROC of 0.9618 (**Figure 7C**) and AUCPR of 0.3559 with the baseline AUCPR of 0.0088 (**Figure 7D**) in the imbalanced dataset, where positive cases (UpDEGs) were significantly outnumbered by negative cases (non-DEGs).

**Figure 7.**
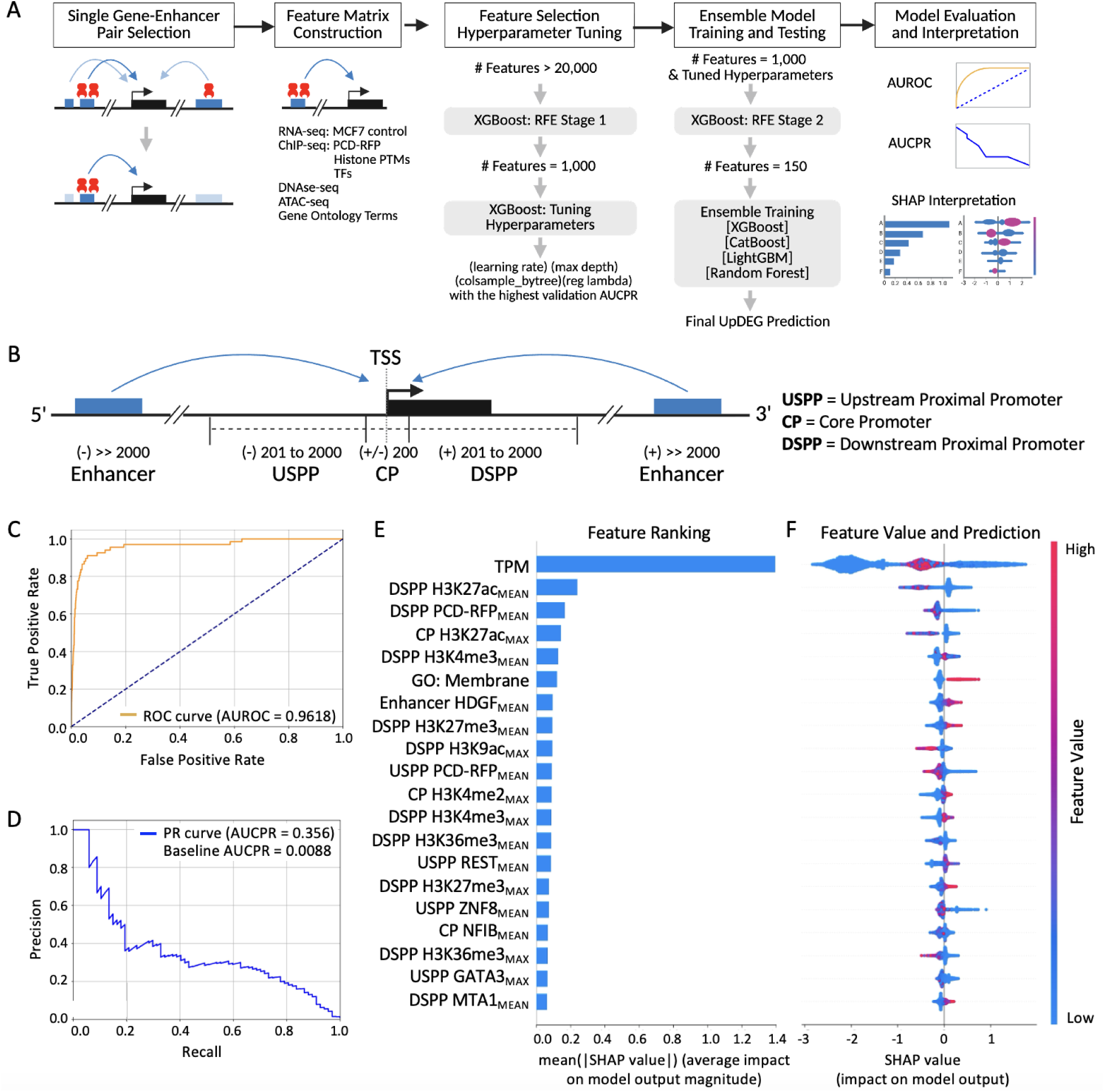
Predictive features at UpDEGs vs. non-DEGs with enhancers that are bound by PCD-RFP. (A) Designation of cis-regulator elements for gene-enhancer pairing at UpDEGs and non-DEGs. (B) Pipeline for machine learning-based prediction of UpDEGs. (C) ROC curve and (D) PR curve for predicting UpDEGs using the filtered 150 features. (E) SHAP bar chart ranking feature importance for UpDEG prediction. (F) SHAP bee swarm chart showing the distribution of feature values in respect to SHAP values.

We used SHAP analysis on the XGBoost component to determine which features were driving the model’s predictions. The baseline gene expression in transcripts per million (TPM) was, by a significant margin, the most important predictive feature (**Figure 7E**). The SHAP bee swarm plot resolved TPM values into three prominent clusters (**Figure 7F**). The two clusters with negative SHAP values suggest that a set of lowly expressed (blue) and highly (red) expressed genes are predicted not to be UpDEGs, while positive UpDEG prediction appeared for a third low-TPM cluster (blue). Together these results suggest that low, not high, TPM is a predictor for UpDEGs, and that not all lowly expressed genes respond to SRA activity. Other features that appear to predict non-DEGs, besides high TPM, are high levels of histone PTMs associated with active chromatin (H3K27ac, H3K9ac, H3K36me3) and transcription factors ZNF8, NFIB, and GATA3 at the promoter regions, which clustered at negative SHAP values.

Interestingly, high PCD-RFP binding at promoter regions was also indicative of non-DEGs. This suggests that TSS-proximal positioning is not optimal for SRA-mediated activation, an idea that is consistent with similar observations from our previous ChIP-seq study in U-2 OS cells.^29^

Features that predict UpDEGs, clustered at positive SHAP values, included H3K4me2 and H3K4me3 at the promoter regions. Taken together with SHAP values that skew towards low baseline TPM, the H3K4me3 signature^51^ suggests that poised or bivalent promoters are a defining feature for UpDEGs. Other predictive promoter features included enrichment of transcription factors REST and MTA1, which are known to function as repressors that interact with histone deacetylases to establish transcriptional silencing.^52–54^ The only enhancer-related element in the top 20 features was the high enrichment of HDGF, a transcription factor that contains a PWWP H3K36me3 reader domain.^55^ Our MLM analyses for PCD-fusion binding also showed high H3K36me3 (**Figure 4**) implicating roles for H3K36me3 in both SRA engagement and subsequent gene activation at UpDEGs. A single gene ontology feature GO:0005886 suggests that genes associated with plasma membrane function are predicted to be UpDEGs. These genes may be epigenetically poised or repressed in MCF7 cells due to their roles in cell adhesion, signaling, and environmental sensing.

## DISCUSSION & CONCLUSIONS

A central goal of targeted epigenetic perturbation is to accurately map the genomic positions that first engage synthetic chromatin regulators and to understand how local chromatin context governs transcriptional outcomes. Our results highlight several best practices for future studies using histone-binding domain (HBD) fusion proteins. First, an HBD-only construct, such as PCD-RFP without an effector, should be used to identify initial contact sites prior to transcriptional activation. Addition of a strong activator could trigger rapid chromatin remodeling that obscures binding events. ChIP-seq performed at substantially earlier time points (30 min - 2 h after induction) could improve resolution of initial interactions. In addition, our findings reinforce the value of topologically associating domains (TADs) as predictive units of gene responsiveness. TADs restrict enhancer-promoter communication and remain conserved across cell types, making them useful landmarks for interpreting SRA-induced gene activation.

Our chromatin feature analyses reveal that a complex chromatin state may be critical for PCD accessibility to H3K27me3-marked nucleosomes. The strong enrichment of additional lysine methylation marks, H3K36me3, H3K4me1, and H4K20me1, at PCD-bound regions suggests an added layer of chromatin information that modulates reader-domain recruitment. These associations are unlikely to be due to biochemical cross-reactivity, as our prior *in vitro* and live-cell data demonstrate high specificity of the CBX8 PCD for H3K27me3.^56,57^ Instead, certain marks might occur on different histone tails within the same nucleosome, or on nearby adjacent nucleosomes, perhaps increasing accessibility of the PCD to H3K27me3.^58^ “Bivalent” single nucleosomes carrying H3K27me3/ H4K20me1, H3K27me3/ H3K36me3, and H3K27me3/ H3K4me1 have been detected in mammalian cells.^51,59^ The co-occurrence of H3K36me3 may reflect recently repressed regions, consistent with the finding that PRC2 binds H3K36me3 prior to depositing H3K27me3.^60^ Our machine-learning analyses do not necessarily suggest physical co-localization of these modifications; rather, they indicate that these features carry predictive value for distinguishing PCD-bound from unbound 200 bp genomic bins. Because our datasets derive from independent, single-PTM ChIP-seq experiments, the spatio-temporal co-occurrence of these marks at PCD target sites remains unresolved and warrants further investigation.

The computational prediction of SRA-responsive genes highlights both the promise and limitations of current machine-learning approaches. To increase precision and recall (e.g., AUCPR values), future analyses will require at least two or three cell lines with distinct epigenomes, genome-wide mapping of PCD-fusion binding, and matched histone PTM and single-cell RNA-seq datasets across early time points. Even earlier profiling (30 min - 4 h) may capture transcription initiation events but could miss delayed responses arising from less accessible chromatin. Our approach used chromosomes 1-17 for training and validating while leaving chromosomes 18-22, X, and Y exclusively for testing, but this partitioning may influence feature clustering profiles. Ideally, a model trained and validated on all chromosomes in one cell line would be tested on all chromosomes from a second, epigenetically distinct line. Accurate enhancer-gene pairing also remains a major limitation. While GeneHancer provides a working approximation, cell-type-specific enhancer-promoter maps would require scHi-C, scATAC-seq, and CRISPR interference perturbations at genome scale, technologies not yet feasible at the required resolution. While ideal datasets remain aspirational, the predictions presented here still provide meaningful insight into SRA-responsive chromatin states.

Across multiple orthogonal analyses, our data converge on a clear conclusion: SRA activity originates at enhancers, not promoters. PCD-bound chromatin states align with enhancer-associated signatures, SRA ChIP-seq peaks are enriched in intronic and distal intergenic regions, and promoter-proximal PCD binding is predictive of nonresponsive genes. Apparent discrepancies in enhancer-feature rankings (e.g., in Fig. 7E) likely reflect the dual mechanism of SRA function, whereby PCD dictates binding at enhancer regions while VP64 activity is constrained by promoter chromatin configuration. Moreover, some UpDEGs may reflect secondary induction rather than direct SRA targeting. Finally, this work advances basic research by demonstrating that SRAs can serve as in situ functional probes for identifying candidate mammalian Polycomb Response Elements (PREs), although further studies in developmental or stem cell contexts may yield clearer overlaps with experimentally validated PREs. Prior efforts relied on laborious reporter assays requiring cloning of candidate PREs upstream of heterologous enhancers, whereas SRAs enable interrogation of regulatory function within native chromatin architecture. Taken together, our findings position SRAs as powerful tools to decode enhancer logic, identify latent regulatory elements, including putative PREs, and dissect the chromatin features that determine whether a silent gene is transcriptionally inducible.

## MATERIALS AND METHODS

### Cell Culture

All cell lines were grown at 37 °C in cell media with high glucose DMEM (Gibco #11965092), 20% tet system approved FBS (Gibco #A4736401), 0.01 mg/mL insulin (Gibco #12585014), and 2 mM L-Glutamine (Gibco #25030081), with 5% CO2 in a humidified incubator. 1X DPBS without calcium and magnesium (Corning #20-031-CV) and 0.25% trypsin-EDTA (Gibco #25200056) was used for cell washing and harvesting. For long-term storage, 1-2 * 10^6 cells were frozen in 1 mL of 10% DMSO (Sigma #D8418) in tet system approved FBS at -80 °C for 2 days, then transferred to -196 °C (liquid nitrogen).

### Transgenic Cell Lines and Transgene Verification

Transgenic MCF7 cell lines were generated as previously described.^31^ To confirm the presence of the transgene, genomic DNA (gDNA) was extracted from transfected cells using the GenElute™ Mammalian Genomic DNA Miniprep Kit (Sigma #G1N70) following the manufacturer’s protocol. PCR was performed with a common primer pair designed to amplify 1469 bp and 1313 bp regions unique to the SRA and PCD-RFP transgenes, respectively (Forward: 5’-[ATCGCCTGGAGCCAATTCC]-3’; Reverse: 5’-[AACCTCCCACATCTCCCCC]-3’). Each 10 µL reaction contained 200 ng of gDNA, 0.5 µM of each primer, and 5 µL of 2X DreamTaq PCR Master Mixes (Thermo #K1081). Thermocycling conditions were as follows: initial denaturation at 95°C for 5 min; 30 cycles of 95°C for 15 s, 52°C for 15 s, and 72°C for 30 s; followed by a final extension at 72°C for 3 min. The resulting amplicons were analyzed by electrophoresis on a 0.8% agarose gel stained with SYBR™ Safe DNA Gel Stain (Invitrogen #S33102). SRA and PCD-RFP transgene inserts were mapped via genome fragment cloning, PCR, and Sanger sequencing as previously described.^32^

### RNA-seq and Data Analysis

Transgenic MCF7 cells were seeded at 1×10^6^ cells/mL in a 6-well plate and grown in media without doxycycline or supplemented with 1.0 μg/mL doxycycline for 10, 24, or 48 h (3 replicate wells per condition). Cells were washed once with warm 1X PBS, lysed in the wells, and RNA was extracted and purified as instructed by the RNeasy Mini kit (Qiagen #74104). Frozen RNA (-80°C) was sent to Novogene for library preparation (polyA-enriched) and deep sequencing (paired-end, 150 bp, Q30 ≥85%). Adapter sequences were removed from sequencing reads using Cutadapt v4.1. Trimmed sequences were aligned to genome build GRCh38.p12 using STAR 2.7.1. Counts of how many reads mapped to each gene were computed using htseq-count v0.11.1. Differential expression analysis was performed using DESeq2 (v1.44.0). Violin plots were generated by ggplot2 (v4.0.0). Upregulated differentially expressed genes (UpDEGs) were identified for each experimental comparison using DESeq2 (v1.44.0),^61^ with thresholds of an adjusted *p*-value (*padj*) ≤ 0.05 and an absolute log2(fold-change) ≥ log2(1.5).

### Reverse Transcription Followed by Quantitative PCR (RT-qPCR)

Total RNA was extracted using RNeasy Mini Kit (Qiagen #74104) from SRA and PCD-RFP transfected MCF7 cells cultured in cell media with 0, 0.001, 0.01, 0.1, 0.5, 1 µg/mL of Dox for 15 h. 1 μg RNA was extracted from two replicates (reps) per sample and used to synthesize cDNA using SuperScript™ IV First-Strand Synthesis System (Invitrogen #18091050). qPCR was performed using PowerUp™ SYBR™ Green Master Mix (Applied Biosystems #A25742), 2 μL 1:100 diluted cDNA, a 760 nM F/R primer mix, total volume 15 μL. Reactions were run on a QuantStudio 6 Flex Real-Time PCR System (Thermo #4484642) as follows: 1 cycle [50°C, 2 min], 1 cycle [95°C, 2 min], 40 cycles [95°C, 15 s; 55–60°C, 15 s; 72°C, 1 min]. Gene targets and respective forward (fwd) and reverse (rev) primers are listed in **Supplemental Table S2**. GAPDH normalized dCt values were calculated as dCt_Target = average Ct_Target - average Ct_GAPDH. Control normalized values (ddCt) were calculated ddCt_Target = (dCt_Target with Dox - average dCt_Target without Dox).

### Chromatin Immunoprecipitation and ChIP-seq

ChIP was performed on transgenic MCF7 cell lines with SRA or with PCD-RFP treated with doxycycline (0, 0.001, 0.01, 0.1, 0.5, or 1.0 µg/mL) for 15 h. Cells were crosslinked with 1% formaldehyde (Thermo #28906) for 5 minutes and quenched, followed by nuclei isolation using the truChIP Chromatin Shearing Kit (Covaris #520154). Chromatin was sheared in microTUBE AFA Fiber Pre-Slit Snap-Cap 6×16mm tubes (Covaris #520045) using a M220 Focused-ultrasonicator (Covaris #M220) for 480 s (Peak Incident Power: 75.0 W, Duty Factor: 5.0%, Cycles/Burst: 200, 7°C). Shearing efficiency to a target size of 200-500 bp was confirmed by 1% agarose gel electrophoresis. Sheared chromatin was pre-cleared with Pierce™ ChIP-Grade Protein A/G Plus Agarose (Thermo #26159). Prior to antibody addition, a 50 µL aliquot was removed from each sample to serve as the ‘input’ control and stored. The remaining chromatin was subsequently immunoprecipitated overnight at 4°C with an mCherry (E5D8F) Rabbit mAb (Cell Signaling Technology #43590S). Immune complexes were captured, washed sequentially with a low salt buffer (20 mM UltraPure™ Tris-HCI (Invitrogen #15568025), 2 mM EDTA (Sigma #03690), 1% Triton™ X-100 (Sigma #648463), 0.1% SDS (Sigma #71736), 150 mM NaCl (Sigma #S3014)) and a high salt buffer (20 mM UltraPure™ Tris-HCI, 2 mM EDTA, 1% Triton™ X-100, 0.1% SDS, 500 mM NaCl), and then eluted in elution buffer (1% SDS, 0.1 M NaHCO3 (Sigma #S8761), 0.1 M NaCl). All samples, including inputs, were reverse-crosslinked overnight at 65°C, treated with Proteinase K (Thermo #17916), and purified using the GenElute PCR Clean-Up Kit (Sigma #NA1020).

ChIP-seq was performed on ChIP and input DNA samples from transgenic MCF7 cell lines with SRA or with PCD-RFP treated with doxycycline (0, 0.1, 0.5, or 1.0 µg/mL) for 15 h. The Kapa HyperPrep kit (Roche) was used to generate ChIP-Seq libraries from 200 pg of ChIP or input DNA. Quantification of the final library was performed using Bioanalyzer (Agilent) and Qubit HS DNA (Thermo Fisher). Libraries were sequenced at PE100 using the Illumina platform (Novaseq 6000 and Nextseq 2000). MCF7 cell lines with PCD-RFP treated with doxycycline (0 or 0.1µg/mL) for 15 h did not pass quality control. ChIP-seq data quality was assessed using FastQC (v0.11.9) for read quality, depth, and sample contamination.^62^ Adapter contamination in SRA samples was removed using Trimmomatic software (v0.39).^63^ Alignment to hg38 was completed using bowtie2 (v2.4.2) and uniquely mapped reads were selected using sambamba (v0.6.8).^64,65^ MACS2 (v2.2.9.1) was used to call peaks. An Irreproducibility Discovery Rate (IDR)(v2.0.3) cutoff of 0.05 was used to filter for peaks present in both replicates.^66^ ChIP-seq broadPeak files were used for downstream computational analyses: MCF7 cells treated with 1.0 µg/mL doxycycline for 15 h, expressing either SRA or PCD-RFP (one replicate each).

### Analysis of Public Bioinformatics Datasets

#### 10 Mb Interval ChIP-seq Peak Heatmaps

The hg38 genome (chr1-22, chrX) was partitioned into contiguous 10 Mb bins. UpDEGs and experimental ChIP-seq peaks were mapped to each 10 Mb bins based on hg38 coordinate overlap. The total number of UpDEGs and ChIP-seq peaks overlapping each bin was counted, and heatmaps were generated using GraphPad (v10.6.1).

#### Z-Score Analysis Bar Charts

Public histone ChIP-seq narrowpeak bed files from ENCODE (H3K4me1, ENCFF991HJA; H3K4me2, ENCFF188VRU; H3K4me3, ENCFF268RXB; H3K9ac, ENCFF348DEB; H3K9me3, ENCFF501UHK; H3K27ac, ENCFF491LQY; H3K27me3, ENCFF669NUD; H3K36me3, ENCFF195FSD; H4K20me1, ENCFF714DEQ) were downloaded^67^ and filtered to retain only standard chromosomes (chr1-22, chrX, and chrY). The genomic complement of experimental ChIP-seq peak files (SRA or PCD-RFP) was generated by subtracting peaks in the experimental ChIP-seq files from the hg38 reference object using GenomicRanges (v1.56.2).^68^ The statistical significance of peak overlaps between each experimental or complement ChIP-seq files (SRA or PCD-RFP) and the public histone ChIP-seq files was determined using the permTest function from the R package regioneR (v1.36.0).^69^ We performed 1,000 permutations, randomizing regions against the hg38 genome assembly. Z-scores were recorded and used to generate bar charts.

#### ChIP-seq Peak Annotation

Genomic annotation of SRA and PCD-RFP ChIP-seq broad peaks was performed using the R package ChIPseeker (v1.40.0).^70^ Peaks were functionally annotated using the hg38 human genome assembly and the org.Hs.eg.db annotation database. The promoter region was defined as +/- 3.0 kb relative to the Transcription Start Site (TSS). The distribution and overlap of peaks across genomic features were visualized using the upsetplot function with the vennpie option enabled.

#### TAD Mapping

Topologically Associating Domain (TAD) boundaries for MCF7 cells were derived from public Hi-C data (GSE66733).^48^ Per-chromosome boundary files (hg19) were processed by calculating the midpoint of the provided ‘start’ and ‘end’ coordinates to define a single boundary position. These boundary coordinates were converted to the hg38 genome build using the liftOver function from the R/Bioconductor package rtracklayer (v1.64.0).^71^ Any boundaries that failed to map during liftover were excluded from further analysis. Finally, contiguous TAD domains were defined based on the sorted hg38 boundary positions with insulation scores ≥ 0.4; each TAD region was set to span from one boundary coordinate to the next consecutive boundary. For each chromosome, the first TAD was defined as starting at base pair 1, and the last TAD was defined as ending at the chromosome terminus. UpDEGs and experimental ChIP-seq peaks were mapped to TADs based on hg38 coordinate overlap. Each TAD was assigned unique identifiers ([chr_Num]-[TAD_Num], e.g. chr1-1).

### Machine Learning Prediction of Genome-Wide PCD-RFP Binding

#### Genomic Binning and Generation of Positive and Negative Sets

The hg38 reference genome was partitioned into contiguous, non-overlapping 200 bp bins based on standard chromosome sizes (chr1-22, chrX, chrY) obtained from the UCSC Genome Browser. Positive regions were defined as 200 bp genomic bins that overlapped with a PCD-RFP peak. A corresponding negative set, 15 times the number of the positive set, with 200 bp random genomic regions excluding all positive 200 bp genomic bins flanked by a 1 kb buffer and also excluding the ENCODE unified blacklist (ENCSR636HFF, hg38) regions was generated.^67^

#### GC-content Matching of the Negative Set

The GC content for all positive regions and candidate negative regions was calculated using the hg38 reference genome. To control for sequence composition bias, a final negative set was generated by matching candidate negative regions to positive regions based on GC content. Regions were binned into 2% GC-content intervals, and negative regions were sampled from each bin (random_state=42) to achieve a 10:1 ratio of negative to positive regions per bin.

#### Histone ChIP-seq Signal Quantification

Mean signal intensities for nine public histone ChIP-seq .bigWig files (H3K4me1, ENCFF763NCP; H3K4me2, ENCFF442RRY; H3K4me3, ENCFF163MXP; H3K9ac, ENCFF327XJC; H3K9me3, ENCFF481DZL; H3K27ac, ENCFF138YNG; H3K27me3, ENCFF163QKN; H3K36me3, ENCFF910BRP; H4K20me1, ENCFF366GLZ) from ENCODE^72^ were quantified within predefined 200 bp genomic bins using pyBigWig (v0.3.24). Genomic intervals with no corresponding data in the source bigWig file, or on chromosomes absent from the file, were assigned a signal value of 0.0.

#### XGBoost Model Training and Evaluation

To prevent data leakage and ensure genomic separation, genomic bins in the positive and the negative sets from chromosomes 1–13 were assigned to the training set, chromosomes 14–17 to the validation set, and chromosomes 18–22, X, and Y to the test set. A gradient-boosted decision tree model was trained using the XGBoost library (v3.0.1)^73^ to classify genomic bins based on one or all nine histone post-translational modification features (H3K4me1, H3K4me2, H3K4me3, H3K9ac, H3K9me3, H3K27ac, H3K27me3, H3K36me3, H4K20me1) against the PCD-RFP binary label. The model was configured with a ‘binary:logistic’ objective, an eta (learning rate) of 0.1, a max_depth of 6, and a seed of 42. Training was performed for up to 1,000 boosting rounds, utilizing the validation set performance (eval_metric: ‘logloss’, ‘auc’, ‘aucpr’) for early stopping with a patience of 50 rounds. This model was then applied to the held-out test set. Performance metrics (logloss, ROC-AUC, AUPRC, F1-score), a file of all test predictions with associated probabilities, a confusion matrix, feature importance scores (weight, gain, cover), ROC curve, and precision-recall curve were generated based on the test set. Feature contributions for the primary XGBoost model were subsequently analyzed using SHAP (SHapley Additive exPlanations) to identify key predictors (v0.49.1).^74^

### Machine Learning Prediction of SRA-Specific UpDEGs

#### Genomic Curation and Annotation of Enhancer-Gene Training Data

To generate training data, the GeneHancer (v5.25) database of enhancer-gene interactions^50^ was first filtered to retain only entries corresponding to genes detected in our RNA-seq dataset, creating a project-specific baseline table. A positive training set was defined by filtering for enhancer elements associated with SRA-specific upregulated differentially expressed genes (UpDEGs). A negative training set was generated by compiling a comprehensive list of all DEGs (both up- and downregulated) and subsequently excluding all enhancers associated with these genes from the baseline table. We then annotated the target gene for each enhancer in both sets using the Ensembl REST API (v15.10)^75^ to programmatically retrieve its chromosome, start position, end position, and strand orientation. The Transcription Start Site (TSS) was operationally defined as the gene_start coordinate for forward-strand genes and the gene_end coordinate for reverse-strand genes. As a final quality control step, any enhancer-gene pair lacking a valid TSS coordinate was removed.

#### Selection of a Single Enhancer-Gene Pair

We generated two key features for all enhancer-gene pairs associated with either upregulated (UpDEG) or non-differentially expressed (Non-DEG) genes. First, we annotated each pair with its maximum signal intensity on the enhancer region by querying the PCD-RFP ChIP-seq signal dataset. Second, we computed the enhancer-TSS distance, defined as the maximum genomic distance from the gene’s transcription start site (TSS) to an enhancer’s start or end coordinate. For each gene, we selected the single associated enhancer with the maximum signal intensity. In the event of a tie, the element with the largest element-TSS distance was chosen as the definitive representative. This filtering step produced the final feature matrices for both the UpDEG and Non-DEG cohorts used for model training.

#### Multi-omic Feature Matrix Construction

For each gene-enhancer pair, we defined four genomic regions relative to the strand-corrected Transcription Start Site (TSS): a Core Promoter (CP; TSS +/- 200 bp), an Upstream Proximal Promoter (USPP; -2000 bp to -201 bp), a Downstream Proximal Promoter (DSPP; +201 bp to +2000 bp), the Enhancer (defined by GeneHancer coordinates). Within these regions, we quantified signals from 9 MCF7 histone PTM ChIP-seq, 116 MCF7 transcription factor ChIP-seq, MCF7 DNase-seq, and MCF7 ATAC-seq datasets from ENCODE^67^ (**Supplemental Table S3**), calculating maximum and mean signal values to generate a primary feature matrix. This chromatin matrix was then supplemented with transcriptomic data by processing MCF7 total RNA-seq (ENCFF721BRA) from ENCODE^67^ to map Transcripts Per Million (TPM) values to each gene via its gene symbol. Finally, to incorporate functional information, we constructed a binary Gene Ontology (GO) matrix and joined it with our dataset, annotating each gene with a one-hot encoded vector of its associated GO terms (PANTHER v.19.0; GO v2025-07-22).^76^

#### Multi-Stage Feature Selection and Hyperparameter Optimization

Positive (upregulated, UpDEG) and negative (non-differentially expressed, Non-DEG) gene sets were partitioned into chromosome-level crossfold training and validation (chr1–17) and test (chr18-22, X, Y) cohorts. This chromosome-based split ensures genomic region independence between the sets used for model development and final evaluation. The feature space was reduced using a two-stage wrapper-based approach. First, an initial Recursive Feature Elimination (RFE) with an XGBoost estimator (100 estimators, 10% step) reduced the full feature set to the top 1,000 predictors (v3.0.1).^73^

A grid search was performed to co-optimize XGBoost hyperparameters (including learning_rate, max_depth, colsample_bytree, and reg_lambda) and the final feature count for the second RFE stage. To maintain chromosomal independence, model performance was assessed using 10-fold GroupKFold cross-validation, with chromosomes serving as distinct groups. The parameter combination yielding the highest mean area under the precision-recall curve (AUCPR) was selected. Second, after hyperparameter tuning, a final RFE pass (50 estimators, 20% step) selected the optimized number of features from this 1,000-feature intermediate set. All RFE procedures were performed on the combined training and validation data with 10-fold GroupKFold cross-validation.

#### Ensemble Model Training, Evaluation, and Interpretation

The optimal number of boosting rounds for the final XGBoost model (v3.0.1)^73^ was determined by 10-fold chromosome-grouped CV using the selected hyperparameters and feature set. The final XGBoost classifier was then trained on the entire combined training and validation dataset using this optimal number of rounds. Concurrently, LightGBM (v4.6.0),^77^ CatBoost (v1.2.8),^78^ and Random Forest (scikit-learn v1.7.2)^79^ classifiers were trained on the same final feature set to generate a diverse model ensemble. The ensemble’s predictive performance was assessed on the held-out test cohort. The final testing used the following hyperparameters (learning_rate = 0.1, max_depth = 8, colsample_bytree = 0.6, reg_lambda = 50.0, n_features = 150). Final probabilities were generated using an unweighted average (soft-voting) of the predictions from all trained models (XGBoost, LightGBM, CatBoost, and Random Forest). Performance was quantified using the area under the receiver operating characteristic (AUC) and precision-recall (AUCPR) curves. Feature contributions for the primary XGBoost model were subsequently analyzed using SHAP (SHapley Additive exPlanations) to identify key predictors (v0.49.1).^74^

## Acknowledgements

The MCF7 ChromHMM 18-state model data was generously provided by J. Ernst. This work was supported by the NIH NCI (R21CA232244 to K.A.H.). Research reported in this publication was supported in part by the Emory Integrated Genomics Core (EIGC; RRID:SCR_023529) of the Winship Cancer Institute of Emory University and NIH/NCI under award number, 2P30CA138292-04, and the Emory Integrated Computational Core (EICC) (RRID:SCR_023525), subsidized by the Emory University School of Medicine. Additional support was provided by the National Center for Georgia Clinical & Translational Science Alliance of the NIH (UL1TR002378). The content is solely the responsibility of the authors and does not necessarily reflect the official views of the National Institutes of Health.

## REFERENCES

1. Baskin, N.L., and Haynes, K.A. (2019). Chromatin engineering offers an opportunity to advance epigenetic cancer therapy. Nat Struct Mol Biol 26, 842–845.

2. Götz, M., and Torres-Padilla, M.-E. (2025). Stem cells as role models for reprogramming and repair. Science 388, eadp2959.

3. Beshnova, D.A., Cherstvy, A.G., Vainshtein, Y., and Teif, V.B. (2014). Regulation of the nucleosome repeat length in vivo by the DNA sequence, protein concentrations and long-range interactions. PLoS Comput Biol 10, e1003698.

4. Calo, E., and Wysocka, J. (2013). Modification of enhancer chromatin: what, how, and why? Mol Cell 49, 825–837.

5. Todd, C.D., Ijaz, J., Torabi, F., Dovgusha, O., Bevan, S., Cracknell, O., Lohoff, T., Clark, S., Argelaguet, R., Pierce, J., et al. (2025). Epigenetic priming of mammalian embryonic enhancer elements coordinates developmental gene networks. Genome Biol 26, 214.

6. Crispatzu, G., Rehimi, R., Pachano, T., Bleckwehl, T., Cruz-Molina, S., Xiao, C., Mahabir, E., Bazzi, H., and Rada-Iglesias, A. (2021). The chromatin, topological and regulatory properties of pluripotency-associated poised enhancers are conserved in vivo. Nat Commun 12, 4344.

7. Maurya, S.S. (2021). Role of Enhancers in Development and Diseases. Epigenomes 5. 10.3390/epigenomes5040021.

8. Heinz, S., Romanoski, C.E., Benner, C., and Glass, C.K. (2015). The selection and function of cell type-specific enhancers. Nat Rev Mol Cell Biol 16, 144–154.

9. Zentner, G.E., Tesar, P.J., and Scacheri, P.C. (2011). Epigenetic signatures distinguish multiple classes of enhancers with distinct cellular functions. Genome Res 21, 1273–1283.

10. Choukrallah, M.-A., Song, S., Rolink, A.G., Burger, L., and Matthias, P. (2015). Enhancer repertoires are reshaped independently of early priming and heterochromatin dynamics during B cell differentiation. Nat Commun 6, 8324.

11. Friedman, M.J., Wagner, T., Lee, H., Rosenfeld, M.G., and Oh, S. (2024). Enhancer-promoter specificity in gene transcription: molecular mechanisms and disease associations. Exp Mol Med 56, 772–787.

12. Creyghton, M.P., Cheng, A.W., Welstead, G.G., Kooistra, T., Carey, B.W., Steine, E.J., Hanna, J., Lodato, M.A., Frampton, G.M., Sharp, P.A., et al. (2010). Histone H3K27ac separates active from poised enhancers and predicts developmental state. Proc Natl Acad Sci U S A 107, 21931–21936.

13. Ramesh, V., Liu, F., Minto, M.S., Chan, U., and West, A.E. (2023). Bidirectional regulation of postmitotic H3K27me3 distributions underlie cerebellar granule neuron maturation dynamics. Elife 12. 10.7554/eLife.86273.

14. Rada-Iglesias, A., Bajpai, R., Swigut, T., Brugmann, S.A., Flynn, R.A., and Wysocka, J. (2011). A unique chromatin signature uncovers early developmental enhancers in humans. Nature 470, 279–283.

15. Tao, L., Yu, H.V., Llamas, J., Trecek, T., Wang, X., Stojanova, Z., Groves, A.K., and Segil, N. (2021). Enhancer decommissioning imposes an epigenetic barrier to sensory hair cell regeneration. Dev Cell 56, 2471–2485.e5.

16. Bleckwehl, T., Crispatzu, G., Schaaf, K., Respuela, P., Bartusel, M., Benson, L., Clark, S.J., Dorighi, K.M., Barral, A., Laugsch, M., et al. (2021). Enhancer-associated H3K4 methylation safeguards in vitro germline competence. Nat Commun 12, 5771.

17. Ostuni, R., Piccolo, V., Barozzi, I., Polletti, S., Termanini, A., Bonifacio, S., Curina, A., Prosperini, E., Ghisletti, S., and Natoli, G. (2013). Latent enhancers activated by stimulation in differentiated cells. Cell 152, 157–171.

18. Bauer, M., Trupke, J., and Ringrose, L. (2016). The quest for mammalian Polycomb response elements: are we there yet? Chromosoma 125, 471–496.

19. Kassis, J.A., and Brown, J.L. (2013). Polycomb group response elements in Drosophila and vertebrates. Adv Genet 81, 83–118.

20. Sing, A., Pannell, D., Karaiskakis, A., Sturgeon, K., Djabali, M., Ellis, J., Lipshitz, H.D., and Cordes, S.P. (2009). A vertebrate Polycomb response element governs segmentation of the posterior hindbrain. Cell 138, 885–897.

21. Woo, C.J., Kharchenko, P.V., Daheron, L., Park, P.J., and Kingston, R.E. (2010). A region of the human HOXD cluster that confers polycomb-group responsiveness. Cell 140, 99–110.

22. Woo, C.J., Kharchenko, P.V., Daheron, L., Park, P.J., and Kingston, R.E. (2013). Variable requirements for DNA-binding proteins at polycomb-dependent repressive regions in human HOX clusters. Mol Cell Biol 33, 3274–3285.

23. Verma, A., Arya, R., and Brahmachari, V. (2022). Identification of a polycomb responsive region in human HoxA cluster and its long-range interaction with polycomb enriched genomic regions. Gene 845, 146832.

24. Cuddapah, S., Roh, T.-Y., Cui, K., Jose, C.C., Fuller, M.T., Zhao, K., and Chen, X. (2012). A novel human polycomb binding site acts as a functional polycomb response element in Drosophila. PLoS One 7, e36365.

25. Metzner, E., Southard, K.M., and Norman, T.M. (2024). Multiome Perturb-seq unlocks scalable discovery of integrated perturbation effects on the transcriptome and epigenome. Cell Syst, 101161.

26. Bracken, A.P., Dietrich, N., Pasini, D., Hansen, K.H., and Helin, K. (2006). Genome-wide mapping of Polycomb target genes unravels their roles in cell fate transitions. Genes Dev. 20, 1123–1136.

27. Hilton, I.B., D’Ippolito, A.M., Vockley, C.M., Thakore, P.I., Crawford, G.E., Reddy, T.E., and Gersbach, C.A. (2015). Epigenome editing by a CRISPR-Cas9-based acetyltransferase activates genes from promoters and enhancers. Nat Biotechnol 33, 510–517.

28. Cai, Y., Zhang, Y., Loh, Y.P., Tng, J.Q., Lim, M.C., Cao, Z., Raju, A., Lieberman Aiden, E., Li, S., Manikandan, L., et al. (2021). H3K27me3-rich genomic regions can function as silencers to repress gene expression via chromatin interactions. Nat Commun 12, 719.

29. Nyer, D.B., Daer, R.M., Vargas, D., Hom, C., and Haynes, K.A. (2017). Regulation of cancer epigenomes with a histone-binding synthetic transcription factor. NPJ Genom Med 2. 10.1038/s41525-016-0002-3.

30. Tillotson, R., Selfridge, J., Koerner, M.V., Gadalla, K.K.E., Guy, J., De Sousa, D., Hector, R.D., Cobb, S.R., and Bird, A. (2017). Radically truncated MeCP2 rescues Rett syndrome-like neurological defects. Nature 550, 398–401.

31. Olney, K.C., Nyer, D.B., Vargas, D.A., Wilson Sayres, M.A., and Haynes, K.A. (2018). The synthetic histone-binding regulator protein PcTF activates interferon genes in breast cancer cells. BMC Syst. Biol. 12, 83.

32. Townsel, A., Wu, Y., Jaffe, M., Shields, C., and Haynes, K.A. (2025). Tet Transgene Activation is Disrupted in Lipogenic Triple Negative Breast Cancer Cells. ACS Synth Biol 14, 2455–2464.

33. Connelly, K.E., Weaver, T.M., Alpsoy, A., Gu, B.X., Musselman, C.A., and Dykhuizen, E.C. (2019). Engagement of DNA and H3K27me3 by the CBX8 chromodomain drives chromatin association. Nucleic Acids Res 47, 2289–2305.

34. Thorvaldsdóttir, H., Robinson, J.T., and Mesirov, J.P. (2013). Integrative Genomics Viewer (IGV): high-performance genomics data visualization and exploration. Brief Bioinform 14, 178–192.

35. Roadmap Epigenomics https://egg2.wustl.edu/roadmap/web_portal/chr_state_learning.html.

36. Ernst, J., Kheradpour, P., Mikkelsen, T.S., Shoresh, N., Ward, L.D., Epstein, C.B., Zhang, X., Wang, L., Issner, R., Coyne, M., et al. (2011). Mapping and analysis of chromatin state dynamics in nine human cell types. Nature 473, 43–49.

37. Roadmap Epigenomics Consortium, Kundaje, A., Meuleman, W., Ernst, J., Bilenky, M., Yen, A., Heravi-Moussavi, A., Kheradpour, P., Zhang, Z., Wang, J., et al. (2015). Integrative analysis of 111 reference human epigenomes. Nature 518, 317–330.

38. Creppe, C., Palau, A., Malinverni, R., Valero, V., and Buschbeck, M. (2014). A Cbx8-containing polycomb complex facilitates the transition to gene activation during ES cell differentiation. PLoS Genet 10, e1004851.

39. Hong, J., Lee, J.H., Zhang, Z., Wu, Y., Yang, M., Liao, Y., de la Rosa, R., Scheirer, J., Pechacek, D., Zhang, N., et al. (2022). PRC2-Mediated Epigenetic Suppression of Type I IFN-STAT2 Signaling Impairs Antitumor Immunity in Luminal Breast Cancer. Cancer Res 82, 4624–4640.

40. Tan, J., Yang, X., Zhuang, L., Jiang, X., Chen, W., Lee, P.L., Karuturi, R.K.M., Tan, P.B.O., Liu, E.T., and Yu, Q. (2007). Pharmacologic disruption of Polycomb-repressive complex 2-mediated gene repression selectively induces apoptosis in cancer cells. Genes Dev. 21, 1050–1063.

41. Lu, Z., Hong, C.C., Kong, G., Assumpção, A.L.F.V., Ong, I.M., Bresnick, E.H., Zhang, J., and Pan, X. (2018). Polycomb Group Protein YY1 Is an Essential Regulator of Hematopoietic Stem Cell Quiescence. Cell Rep 22, 1545–1559.

42. Konuma, T., Oguro, H., and Iwama, A. (2010). Role of the polycomb group proteins in hematopoietic stem cells. Dev Growth Differ 52, 505–516.

43. Schoenfelder, S., Sugar, R., Dimond, A., Javierre, B.-M., Armstrong, H., Mifsud, B., Dimitrova, E., Matheson, L., Tavares-Cadete, F., Furlan-Magaril, M., et al. (2015). Polycomb repressive complex PRC1 spatially constrains the mouse embryonic stem cell genome. Nat Genet 47, 1179–1186.

44. Szabo, Q., Bantignies, F., and Cavalli, G. (2019). Principles of genome folding into topologically associating domains. Sci Adv 5, eaaw1668.

45. Dixon, J.R., Gorkin, D.U., and Ren, B. (2016). Chromatin Domains: The Unit of Chromosome Organization. Mol Cell 62, 668–680.

46. Qiao, Y., Wang, Z., Tan, F., Chen, J., Lin, J., Yang, J., Li, H., Wang, X., Sali, A., Zhang, L., et al. (2020). Enhancer Reprogramming within Pre-existing Topologically Associated Domains Promotes TGF-β-Induced EMT and Cancer Metastasis. Mol Ther 28, 2083–2095.

47. McArthur, E., and Capra, J.A. (2021). Topologically associating domain boundaries that are stable across diverse cell types are evolutionarily constrained and enriched for heritability. Am J Hum Genet 108, 269–283.

48. Barutcu, A.R., Lajoie, B.R., McCord, R.P., Tye, C.E., Hong, D., Messier, T.L., Browne, G., van Wijnen, A.J., Lian, J.B., Stein, J.L., et al. (2015). Chromatin interaction analysis reveals changes in small chromosome and telomere clustering between epithelial and breast cancer cells. Genome Biol 16, 214.

49. Williamson, I., Graham, K.A., Woolf, M., Becher, H., Hill, R.E., Bickmore, W.A., and Lettice, L.A. (2025). Bystander activation across a TAD boundary supports a cohesin-dependent transcription cluster model for enhancer function. Genes Dev 39, 1012–1024.

50. Fishilevich, S., Nudel, R., Rappaport, N., Hadar, R., Plaschkes, I., Iny Stein, T., Rosen, N., Kohn, A., Twik, M., Safran, M., et al. (2017). GeneHancer: genome-wide integration of enhancers and target genes in GeneCards. Database (Oxford) 2017. 10.1093/database/bax028.

51. Voigt, P., LeRoy, G., Drury, W.J., 3rd, Zee, B.M., Son, J., Beck, D.B., Young, N.L., Garcia, B.A., and Reinberg, D. (2012). Asymmetrically modified nucleosomes. Cell 151, 181–193.

52. Thiel, G., Ekici, M., and Rössler, O.G. (2015). RE-1 silencing transcription factor (REST): a regulator of neuronal development and neuronal/endocrine function. Cell Tissue Res 359, 99–109.

53. Nassar, A., Satarker, S., Gurram, P.C., Upadhya, D., Fayaz, S.M., and Nampoothiri, M. (2023). Repressor Element-1 Binding Transcription Factor (REST) as a Possible Epigenetic Regulator of Neurodegeneration and MicroRNA-Based Therapeutic Strategies. Mol Neurobiol 60, 5557–5577.

54. Molli, P.R., Singh, R.R., Lee, S.W., and Kumar, R. (2008). MTA1-mediated transcriptional repression of BRCA1 tumor suppressor gene. Oncogene 27, 1971–1980.

55. Wang, H., Farnung, L., Dienemann, C., and Cramer, P. (2020). Structure of H3K36-methylated nucleosome-PWWP complex reveals multivalent cross-gyre binding. Nat Struct Mol Biol 27, 8–13.

56. Franklin, K.A., Priode, J.H., Steppe, P., and Haynes, K.A. (2024). An engineered chromatin protein with enhanced preferential binding of H3K27me3 over H3K9me3. bioRxiv. 10.1101/2024.10.02.616304.

57. Tekel, S.J., Vargas, D.A., Song, L., LaBaer, J., Caplan, M.R., and Haynes, K.A. (2018). Tandem Histone-Binding Domains Enhance the Activity of a Synthetic Chromatin Effector. ACS Synth Biol 7, 842–852.

58. Bryan, E., Valsakumar, D., Idigo, N.J., Warburton, M., Webb, K.M., McLaughlin, K.A., Spanos, C., Lenci, S., Major, V., Ambrosi, C., et al. (2025). Nucleosomal asymmetry shapes histone mark binding and promotes poising at bivalent domains. Mol Cell 85, 471–489.e12.

59. Chen, Z., Song, L., Chen, W., and Wang, C. (2025). Single-cell profiling of H3K4me1-H3K27me3 revealed bivalent regulation of abnormal neuronal development caused by prenatal e-cigarette vaporing. Commun Biol 8, 1326.

60. Brien, G.L., Gambero, G., O’Connell, D.J., Jerman, E., Turner, S.A., Egan, C.M., Dunne, E.J., Jurgens, M.C., Wynne, K., Piao, L., et al. (2012). Polycomb PHF19 binds H3K36me3 and recruits PRC2 and demethylase NO66 to embryonic stem cell genes during differentiation. Nat Struct Mol Biol 19, 1273–1281.

61. Love, M.I., Huber, W., and Anders, S. (2014). Moderated estimation of fold change and dispersion for RNA-seq data with DESeq2. Genome Biol 15, 550.

62. 62. Website http://www.bioinformatics.babraham.ac.uk/projects/fastqc.

63. Bolger, A.M., Lohse, M., and Usadel, B. (2014). Trimmomatic: a flexible trimmer for Illumina sequence data. Bioinformatics 30, 2114–2120.

64. Langmead, B., and Salzberg, S.L. (2012). Fast gapped-read alignment with Bowtie 2. Nat. Methods 9, 357–359.

65. Zhang, Y., Liu, T., Meyer, C.A., Eeckhoute, J., Johnson, D.S., Bernstein, B.E., Nusbaum, C., Myers, R.M., Brown, M., Li, W., et al. (2008). Model-based analysis of ChIP-Seq (MACS). Genome Biol 9, R137.

66. Li, Q., Brown, J.B., Huang, H., and Bickel, P.J. (2011). Measuring reproducibility of high-throughput experiments. Ann. Appl. Stat. 5, 1752–1779.

67. Luo, Y., Hitz, B.C., Gabdank, I., Hilton, J.A., Kagda, M.S., Lam, B., Myers, Z., Sud, P., Jou, J., Lin, K., et al. (2020). New developments on the Encyclopedia of DNA Elements (ENCODE) data portal. Nucleic Acids Res 48, D882–D889.

68. Lawrence, M., Huber, W., Pagès, H., Aboyoun, P., Carlson, M., Gentleman, R., Morgan, M.T., and Carey, V.J. (2013). Software for computing and annotating genomic ranges. PLoS Comput Biol 9, e1003118.

69. Gel, B., Díez-Villanueva, A., Serra, E., Buschbeck, M., Peinado, M.A., and Malinverni, R. (2016). regioneR: an R/Bioconductor package for the association analysis of genomic regions based on permutation tests. Bioinformatics 32, 289–291.

70. Wang, Q., Li, M., Wu, T., Zhan, L., Li, L., Chen, M., Xie, W., Xie, Z., Hu, E., Xu, S., et al. (2022). Exploring Epigenomic Datasets by ChIPseeker. Curr Protoc 2, e585.

71. Lawrence, M., Gentleman, R., and Carey, V. (2009). rtracklayer: an R package for interfacing with genome browsers. Bioinformatics 25, 1841–1842.

72. Ryan, D., Roberts, E., Eraslan, G., Grüning, B., Ward, D., Jamous, B.A., Betts, E., Ramirez, F., Fox, N., Abdennur, N., et al. deeptools/pyBigWig: 0.3.24. 10.5281/zenodo.14679776.

73. XGBoost 10.1145/2939672.2939785.

74. Lundberg, S.M., Erion, G., Chen, H., DeGrave, A., Prutkin, J.M., Nair, B., Katz, R., Himmelfarb, J., Bansal, N., and Lee, S.-I. (2020). From local explanations to global understanding with explainable AI for trees. Nature Machine Intelligence 2, 56–67.

75. Yates, A., Beal, K., Keenan, S., McLaren, W., Pignatelli, M., Ritchie, G.R.S., Ruffier, M., Taylor, K., Vullo, A., and Flicek, P. (2015). The Ensembl REST API: Ensembl Data for Any Language. Bioinformatics 31, 143–145.

76. Thomas, P.D., Ebert, D., Muruganujan, A., Mushayahama, T., Albou, L.-P., and Mi, H. (2022). PANTHER: Making genome-scale phylogenetics accessible to all. Protein Sci. 31, 8–22.

77. Ke, G., Meng, Q., Finley, T., Wang, T., Chen, W., Ma, W., Ye, Q., and Liu, T.-Y. (2017). LightGBM: A Highly Efficient Gradient Boosting Decision Tree. In Neural Information Processing Systems, pp. 3149–3157.

78. Dorogush, A.V., Ershov, V., and Gulin, A. (2018). CatBoost: gradient boosting with categorical features support.

79. Pedregosa, F., Varoquaux, G., Gramfort, A., Michel, V., Thirion, B., Grisel, O., Blondel, M., Prettenhofer, P., Weiss, R., Dubourg, V., et al. (2011). Scikit-learn: Machine Learning in Python. J. Mach. Learn. Res. 12, 2825–2830.

