## Supplemental Information for "Decoding Gene Responsiveness to Synthetic Chromatin Reader-Actuators with Multi-Modal Epigenomic Profiling"

**Supplemental Figures**

1. Figure S1. Comparison of histone PTM overlaps with PCD-RFP and SRA ChIP-seq peaks with ChromHMM regulatory states
2. Figure S2. ChIP-Atlas fold enrichment analysis of histone PTM at fusion protein ChIP-seq peaks

**Supplemental Tables**

1. Table S1. Transcription factor target genes within SRA UpDEG classes
2. Table S2. RT-qPCR Primer Pairs
3. Table S3. ENCODE MCF7 Datasets

**Supplemental Materials and Methods**

1. Analysis of Public Bioinformatics Datasets (for Figure S1)

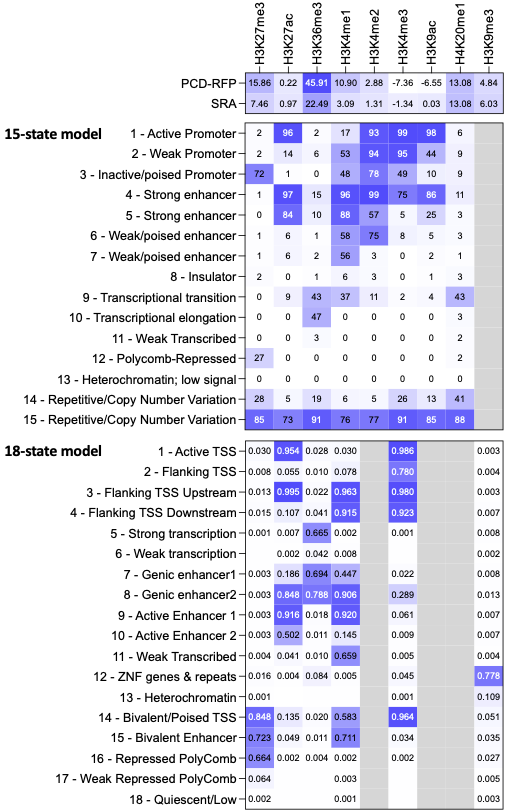

**Figure S1. Comparison of histone PTM overlaps with PCD-RFP and SRA ChIP-seq peaks with ChromHMM regulatory states.** Top table: Z-score values of histone PTM ChIP-seq peaks and PCD-RFP or SRA ChIP-seq peak overlaps. Middle table: 15-state ChromHMM model with chromatin mark observation frequencies from Figure 1 in J. Ernst et al.^1^ Bottom table: 18-state Chrom HMM model from the Epigenome Roadmap (https://egg2.wustl.edu/roadmap/web_portal/chr_state_learning.html).

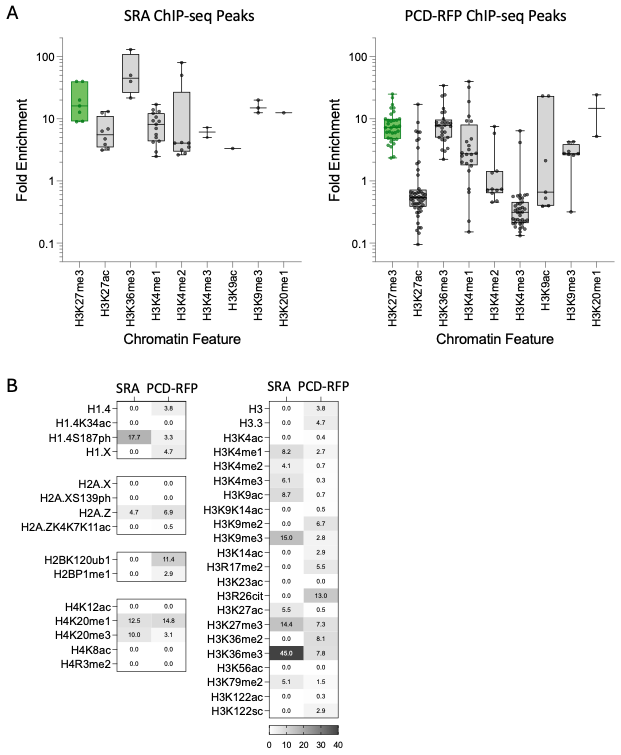

**Figure S2. ChIP-Atlas fold enrichment analysis of histone PTM at fusion protein ChIP-seq peaks.** (A) Fold enrichment (FE) of chromatin features within ChIP-seq peak regions for SRA and PCD-RFP versus randomly permuted regions. Box plots include data for regulatory state-associated histone PTMs H3K27me3, H3K27ac, H3K36me3, H3K4me1, H3K4me2, H3K4me3, H3K9ac, H3K9me3, and H3K20me1. Only statistically significant values (*p* < 0.001) are included. (B) Tables show median enrichments for all histone modifications that showed at least one significant FE value.

**SUPPLEMENTAL TABLES**

**Table S1. Transcription factor target genes within SRA UpDEG classes**

| **TF** | **FC** | **Target gene UpDEGs** | | | | | |
| --- | --- | --- | --- | --- | --- | --- | --- |
|  |  | Early transient | Early sustained | Late transient | Late sustained | Delayed | Stochastic |
| HOXB6  (UpDEG) | 0.60 | (11) CSRP2, EYA4, HOXC9, ISM2, LRRC8C, MAFA, MFAP3L, NPL, PPP4R4, SRPX, TFPI2 | (5) BOC, EPHB2, FAM171A1, FLT1, SEMA6B | (5) ARHGDIG, BACH2, ETV4, PANX2, ZMYND15 | (3) COLEC12, MAPRE2, PDIA2 | (6) ABCA10, ADD3, DMD, FAT4, TECTA, WIPF1 | none |
| HOXB9  (UpDEG) | 0.83 | (16) CSRP2, DDR2, DLX3, FGD1, GPR156, IGF2BP1, ISM1, KBTBD11, KHDRBS3, MFAP3L, NPL, OLFML2A, PARVB, PLSCR4, PRR5L, SMAD9 | (6) EPHB2, ESRRG, FLT1, LIMD2, LIPG, SEMA6B | (5) ETV4, ETV5, F2RL1, PNPLA3, TMEM169 | (4) MAPRE2, PNCK, SPRY4, TIMP2 | (10) DISC1, IGF2, LTBP3, MMP24, NR2F1, RBMS2, RPRM, SEMA6A, TMPRSS11E, TNIK | (1) CHSY3 |
| HOXB13  (UpDEG) | 1.02 | (22) CLIP2, ERG, HOXC9, HTR6, IGF2BP1, ISM1, KBTBD11, KCNG1, KHDRBS3, KIF21B, MFAP3L, PLCL2, PLSCR4, PRR5L, RGMA, RGS9, RIMS3, STEAP2, SYNE3, TNFAIP8L3, TRNP1, TSHZ3 | (4) EPHB2, GJB2, HTR1D, SEMA6B | (6) ARHGDIG, ANTXR2, F2RL1, GXYLT2, TMEM169, WFDC2 | (2) PDIA2, SPRY4 | (8) RBMS2, RPRM, SEMA6A, SLC25A27, STON1, TMPRSS11E, TNIK, WIPF1 | (1) LRP12 |
| HOXC8  (UpDEG) | 1.08 | (12) BCL2L10, CSRP2, DDR2, FAM78A, HOXC9, ISM2, MAFA, NPL, NRTN, PPP4R4, STEAP2, TRNP1 | (5) ESRRG, FAM171A1, GMPR, RASGRP2, SEMA6B | (5) BACH2, ETV4, PDLIM3, PIK3AP1, ZMYND15 | (2) DPP4, MAPRE2 | (4) ADD3, SEMA6A, SMPD1, WIPF1 | (1) JAZF1 |
| HOXC9  (UpDEG) | 0.64 | (10) CSRP2, DPYSL5, FAM78A, HOXC9, KBTBD11, KHDRBS3, KIAA1549L, NRTN, PRR5L, RCAN2 | (2) ESRRG, FLT1 | (3) BACH2, ETV4, THRA | (2) BEND7, MAPRE2 | (5) DISC1, FAT4, MMP24, SEMA6A, SEMA6D | (1) JAZF1 |
| HOXC10  (UpDEG) | 0.64 | (7) DPYSL5, FAM78A, HOXC8, KHDRBS3, KIF21B, RYR1, TUBB2B | (4) ESRRG, FLT1, KIF1A, SERPINE2 | (3) GXYLT2, PNPLA3, SLC29A4 | (2) BEND7, FBLN7 | (3) DISC1, DMD, STON1 | (1) JAZF1 |
| YY1 | n/a | (14) BCL2L10, DDR2, DPYSL5, GLIPR2, HAP1, HOXC10, KCNG1, MFAP3L, PRIMA1, PRR5L, RASSF5, STEAP2, TNFAIP8L3, TUBB2B | (8) CYBRD1, FXYD6, GJB2, GMPR, HTR1D, LARP6, LIPG, RASL10B | (4) ETV4, GXYLT2, PDLIM3, TMEM86A | (5) ATP1B2, BEAN1, COL23A1, COLEC12, DBN1 | (4) ADD3, IGF2, STON1, WIPF1 | (1) LGI2 |

The TF Database^2^ was used to retrieve all target promoters and genes for each transcription factor (TF) shown in the first column. Target gene UpDEGs were identified by cross-checking this output with the entire list of 397 UpDEGs. For TFs that are also UpDEGs, the maximum log2 fold change (FC) across the three time points (10, 24, 48 hours) is shown in the second column.

**Table S2. RT-qPCR Primer Pairs**

| **Target** | **Forward 5’ to 3’** | **Reverse 5’ to 3’** |
| --- | --- | --- |
| SRA | ATCGCCTGGAGCCAATTCC | AACCTCCCACATCTCCCCC |
| PCD-RFP | ATCGCCTGGAGCCAATTCC | AACCTCCCACATCTCCCCC |
| GAPDH | TCTCCTCTGACTTCAACAGCGAC | CCCTGTTGCTGTAGCCAAATTC |

The same primer pair was used for the fusion protein transcripts, SRA and PCD-RFP, which binds to the mCherry sequence that is present in both mRNAsequences.

**Table S3. ENCODE MCF7 Datasets**

| **Type** | **Feature** | **ENCODE Accession** |
| --- | --- | --- |
| Histone ChIP-seq | H3K4me1 | ENCFF763NCP |
|  | H3K4me2 | ENCFF442RRY |
|  | H3K4me3 | ENCFF163MXP |
|  | H3K9ac | ENCFF327XJC |
|  | H3K9me3 | ENCFF481DZL |
|  | H3K27ac | ENCFF138YNG |
|  | H3K27me3 | ENCFF163QKN |
|  | H3K36me3 | ENCFF910BRP |
|  | H4K20me1 | ENCFF366GLZ |
| Transcription Factor ChIP-seq | ARID3A | ENCFF907DWT |
|  | ATF7 | ENCFF703ZRP |
|  | BMI1 | ENCFF977TJK |
|  | CEBPB | ENCFF464JOC |
|  | CHD1 | ENCFF923MDX |
|  | CLOCK | ENCFF045RXD |
|  | COPS2 | ENCFF102ABZ |
|  | CREB1 | ENCFF954WHX |
|  | CSDE1 | ENCFF722GUE |
|  | CTBP1 | ENCFF922FKY |
|  | CTCF | ENCFF507CRU |
|  | CUX1 | ENCFF663YXL |
|  | DDX20 | ENCFF895IXP |
|  | DPF2 | ENCFF142KHL |
|  | E2F8 | ENCFF919YCR |
|  | E4F1 | ENCFF555HAI |
|  | EGR1 | ENCFF487DKA |
|  | ELF1 | ENCFF751TAA |
|  | ELK1 | ENCFF045XJM |
|  | EP300 | ENCFF708NMR |
|  | ESRRA | ENCFF059ZSC |
|  | FOS | ENCFF327LTA |
|  | FOSL2 | ENCFF704HAX |
|  | FOXA1 | ENCFF512UGW |
|  | FOXK2 | ENCFF477VZD |
|  | FOXM1 | ENCFF405RBL |
|  | GABPA | ENCFF494GAR |
|  | GATA3 | ENCFF971AZB |
|  | GATAD2B | ENCFF560WGB |
|  | GTF2F1 | ENCFF736KUY |
|  | HCFC1 | ENCFF103ABY |
|  | HDAC2 | ENCFF178WAQ |
|  | HDGF | ENCFF178LPH |
|  | HES1 | ENCFF509HTJ |
|  | HSF1 | ENCFF505ZBM |
|  | JUN | ENCFF730TVS |
|  | JUND | ENCFF990FGN |
|  | LARP7 | ENCFF148SFF |
|  | MAFK | ENCFF174HAP |
|  | MAX | ENCFF315FIO |
|  | MAZ | ENCFF297WGG |
|  | MBD2 | ENCFF517UQD |
|  | MLLT1 | ENCFF654ISN |
|  | MNT | ENCFF623IRB |
|  | MTA1 | ENCFF883HBF |
|  | MTA2 | ENCFF557XGU |
|  | MTA3 | ENCFF486PVE |
|  | NBN | ENCFF242QKO |
|  | NCOA3 | ENCFF770MLI |
|  | NEUROD1 | ENCFF501HJF |
|  | NFIB | ENCFF631HNZ |
|  | NFRKB | ENCFF411WFM |
|  | NFXL1 | ENCFF537TBX |
|  | NONO | ENCFF875CIS |
|  | NR2F2 | ENCFF678MPN |
|  | NRF1 | ENCFF113MFU |
|  | PAX8 | ENCFF775POG |
|  | PKNOX1 | ENCFF737NUU |
|  | PML | ENCFF911FVQ |
|  | POLR2A | ENCFF827YIP |
|  | PPP1R10 | ENCFF167JMR |
|  | RAD21 | ENCFF775EKJ |
|  | RAD51 | ENCFF522WFI |
|  | RCOR1 | ENCFF536KZN |
|  | REST | ENCFF068TEY |
|  | RFX1 | ENCFF310ZPD |
|  | RFX5 | ENCFF085XIV |
|  | SIN3A | ENCFF807QKK |
|  | SIX4 | ENCFF032MSQ |
|  | SMARCA5 | ENCFF297JNO |
|  | SMARCE1 | ENCFF375LRP |
|  | SNIP1 | ENCFF231LYV |
|  | SP1 | ENCFF367HVY |
|  | SREBF1 | ENCFF791DFW |
|  | SRF | ENCFF534AJF |
|  | SUZ12 | ENCFF598JRC |
|  | TAF1 | ENCFF418WWV |
|  | TARDBP | ENCFF065RQM |
|  | TCF7L2 | ENCFF711CKB |
|  | TCF12 | ENCFF031XVC |
|  | TEAD4 | ENCFF977ZGZ |
|  | TOE1 | ENCFF318YIK |
|  | TRIM22 | ENCFF100OCC |
|  | YBX1 | ENCFF033VON |
|  | ZBTB1 | ENCFF062ZHZ |
|  | ZBTB7B | ENCFF073COQ |
|  | ZBTB11 | ENCFF510SQD |
|  | ZBTB33 | ENCFF506CTQ |
|  | ZBTB40 | ENCFF539ZHK |
|  | ZFX | ENCFF482RYD |
|  | ZHX2 | ENCFF724GPW |
|  | ZKSCAN1 | ENCFF239BWH |
|  | ZNF8 | ENCFF550FAH |
|  | ZNF24 | ENCFF505ORH |
|  | ZNF207 | ENCFF668FJE |
|  | ZNF217 | ENCFF491CXP |
|  | ZNF217 | ENCFF343WRL |
|  | ZNF444 | ENCFF101ION |
|  | ZNF507 | ENCFF659LTF |
|  | ZNF512B | ENCFF413BLF |
|  | ZNF574 | ENCFF914VHT |
|  | ZNF579 | ENCFF913XOW |
|  | ZNF592 | ENCFF950SDY |
|  | ZNF687 | ENCFF910XUA |
| Chromatin accessibility | DNase-seq | ENCFF137FTO |
|  | ATAC-seq | ENCFF976UNK |

**SUPPLEMENTAL MATERIALS AND METHODS**

**Analysis of ChIP-Atlas Datasets**

To generate the data shown in Figure S1, the “ChIP-Atlas: Enrichment Analysis” tool (https://chip-atlas.org/enrichment_analysis) was used to determine Fold Enrichment of chromatin features, mapped in MCF7 untreated parental cells, overlapping with regions of interest (Dataset A): SRA ChIP-seq peaks (295), or PCD-RFP ChIP-seq peaks (13477). Settings were as follows: genome, *H. sapiens* (hg38); experiment type, ChIP: Histone (null); cell type class, Breast (2821); threshold for significance = 50; Dataset B, random permutation of Dataset A (x10). P-values were calculated by Fisher's exact test to compare the probability of Dataset A versus B overlaps. Infinite fold enrichment values (where Dataset B overlaps = 0) were omitted. Fold enrichment outputs from drug or hormone treated or genetically manipulated MCF7 cells were excluded from downstream analyses.

**SUPPLEMENTAL REFERENCES**

1.	Ernst, J., Kheradpour, P., Mikkelsen, T.S., Shoresh, N., Ward, L.D., Epstein, C.B., Zhang, X., Wang, L., Issner, R., Coyne, M., et al. (2011). Mapping and analysis of chromatin state dynamics in nine human cell types. Nature *473*, 43–49.

2.	Plaisier, C.L., O’Brien, S., Bernard, B., Reynolds, S., Simon, Z., Toledo, C.M., Ding, Y., Reiss, D.J., Paddison, P.J., and Baliga, N.S. (2016). Causal Mechanistic Regulatory Network for Glioblastoma Deciphered Using Systems Genetics Network Analysis. Cell Syst *3*, 172–186.
